# Functional maturation of human neural stem cells in a 3D bioengineered brain model enriched with fetal brain-derived matrix

**DOI:** 10.1101/691907

**Authors:** Disha Sood, Dana M. Cairns, Jayanth M. Dabbi, Charu Ramakrishnan, Karl Deisseroth, Lauren D. Black, Sabato Santaniello, David L. Kaplan

## Abstract

Brain extracellular matrix (ECM) is often overlooked *in vitro* brain tissue models, despite its instructive roles during development. Using developmental stage-sourced brain ECM in reproducible 3D bioengineered culture systems, we demonstrate enhanced functional differentiation of human induced neural stem cells (hiNSCs) into healthy neurons and astrocytes. Particularly, fetal brain tissue-derived ECM supported long-term maintenance of differentiated neurons, demonstrated by morphology, gene expression and secretome profiling. Astrocytes were evident within the second month of differentiation, and reactive astrogliosis was inhibited in brain ECM-enriched cultures when compared to unsupplemented cultures. Functional maturation of the differentiated hiNSCs within fetal ECM-enriched cultures was confirmed by calcium signaling and unsupervised cluster analysis. Additionally, the study identified native biochemical cues in decellularized ECM with notable comparisons between fetal and adult brain-derived ECMs. The development of novel brain-specific biomaterials for generating mature *in vitro* brain models provides an important path forward for interrogation of neuron-glia interactions.

## Introduction

Many brain physiological and pathological features are human-specific, making it difficult to extrapolate results from animal models, and thus driving the need for innovative human cell-based 3D *in vitro* brain tissue cultures to investigate neurological disorders. Additionally, many neurodevelopmental and neurodegenerative disorders are polygenic with multiple syndromic and non-syndromic forms, some of which are of unknown genetic etiology and thus challenging to investigate using animal models (Avior, Sagi, & Benvenisty, 2016). Previous *in vitro* models of neurological disorders using human induced pluripotent stem cells (hiPSCs) or human neural stem cells (hNSCs) have mainly involved monolayer 2D cultures that do not recapitulate physiological cell phenotype, signaling, and drug sensitivity due to the lack of high cell densities, connectivity in 3D networks, and relevant cell-cell and cell-ECM interactions (de la Torre-Ubieta, Won, Stein, & Geschwind, 2016; Quadrato, Brown, & Arlotta, 2016). Recent advances in 3D organoid and spheroid-based systems have been extremely useful for studies of normal brain development, such as cortical layering/interneuron migration, and for neurodevelopmental disorders such as microcephaly, lissencephaly, and autism (Bagley, Reumann, Bian, Levi-Strauss, & Knoblich, 2017; Birey et al., 2017; Camp et al., 2015; Giandomenico & Lancaster, 2017; Lancaster et al., 2013; Luo et al., 2016; Mariani et al., 2015; A. M. Pasca et al., 2015; Quadrato et al., 2017; Sloan et al., 2017). Despite the variety of approaches there are only a few examples of 3D *in vitro* brain-like tissue models that exhibit neuronal maturity and co-differentiation into glial cell types (Marton et al., 2019; A. M. Pasca et al., 2015; Sloan et al., 2017). Also, most of these models were associated with significant limitations including slow maturation into neuronal supporting cell types, such as astrocytes, and/or necrosis at longer time points of cultivation *in vitro* (Quadrato et al., 2016; Velasco et al., 2019). This maturation and cell-cell interactions are key for modeling synaptogenesis and functions during later postnatal developmental stages and for revealing the molecular basis for many diseased states, particularly neurodegenerative disorders where neuron-glia interactions are dysfunctional (Y. H. Kim et al., 2015; Lian & Zheng, 2016; Rama Rao & Kielian, 2015; Salmina, 2009). Many of these 3D brain-like tissue models are also limited in terms of reproducibility, and compartmentalization related to the introduction of microglia and vasculature, as well as for sampling and control of nutrient transport into and out of the tissue systems.

A common limitation of current 3D *in vitro* brain models is that the ECM content is often not considered in detail, even though brain ECM is dynamic during development and plays a crucial role in cell signaling and homeostasis (Zimmermann & Dours-Zimmermann, 2008). The ‘dynamic reciprocity’ model was proposed in the 1980s, which suggested that ECM guides gene expression and individual components of ECM have an instructive role in directing tissue-specific development (Bissell, Hall, & Parry, 1982). Despite these roles, most 3D brain tissue models use Matrigel as the major ECM component and/or soluble bioactive factors to induce differentiation. Matrigel is a mouse sarcoma-derived basement membrane matrix that lacks many physiologically-relevant biochemical cues involved in brain development and maintenance, including several glycoproteins and proteogylcans (Bandtlow & Zimmermann, 2000; Hughes, Postovit, & Lajoie, 2010; Miyata & Kitagawa, 2017). The human brain ECM constitutes about 20-40% of the brain volume during development and adulthood, is highly organized, and has unique traits in composition when compared to the ECM of other tissues (Zimmermann & Dours-Zimmermann, 2008). Moreover, during development, ECM guides the compartmentalization of functional brain microdomains, and thus contributes to the sophisticated architecture and function of the brain (Dityatev, Seidenbecher, & Schachner, 2010). Such native ECM signals are particularly important for differentiation and complete maturation of neural progenitor/stem cells (Hoshiba et al., 2016).

The impact of adult brain-derived ECM on cell differentiation, gelation kinetics and mechanical properties has been studied in isolation (De Waele et al., 2015; DeQuach et al., 2010; Medberry et al., 2013); however, the study of composite, scaffold-based 3D *in vitro* systems to investigate the bioactivity of ECM from different developmental stages over long-term differentiation of human induced neural stem cells (hiNSCs) into both mature neurons and astrocytes is lacking. Astrocytes respond to soluble factors and also influence their environment through the secretion of ECM molecules, particularly chondroitin sulfate proteoglycans (CSPGs) that vary with mature/resting versus reactive astrocytes (Avram, Shaposhnikov, Buiu, & Mernea, 2014; Lian & Zheng, 2016; Wiese, Karus, & Faissner, 2012). Therefore, preventing reactive astrogliosis, measured by consistently high CSPG release, in 3D *in vitro* brain models is critical in order to maintain neuronal health and functional synapses (Krencik, van Asperen, & Ullian, 2017; Yu, Wang, Katagiri, & Geller, 2012).

We hypothesized that the use of native brain-derived ECM for brain-relevant biochemical cues, in combination with a tissue engineered approach to design brain-specific tissue constructs would promote improved differentiation of stem cells; as well as address the need for reproducibility, tunability for compartmentalization/sampling and accelerated maturation of the cells into neurons and glia. Many ECM proteins are conserved across species (Hynes, 2012), thus porcine brain-derived ECM was used towards the differentiation of hNSCs. In the current study, we investigated the effects of brain-derived ECM from two different developmental stages (fetal versus adult) on the differentiation of hiNSCs (Cairns et al., 2016) into mature neurons and healthy astrocytes, when cultured within relevant environmental cues (biochemical factors and 3D topology) (Schematic 1). hiNSCs were cultured in bioengineered silk protein scaffold-based 3D tissue constructs infused with either collagen type I (CLG1, previously shown to be compatible with brain cells (Chwalek, Tang-Schomer, Omenetto, & Kaplan, 2015)) or hyaluronic acid (HA, a brain ECM component (Charles, Holland, Gilbertson, Glass, & Kettenmann, 2012; Wiranowska & Rojiani, 2011)) hydrogels supplemented with native brain-derived ECM (fetal or adult).

**Schematic 1:**
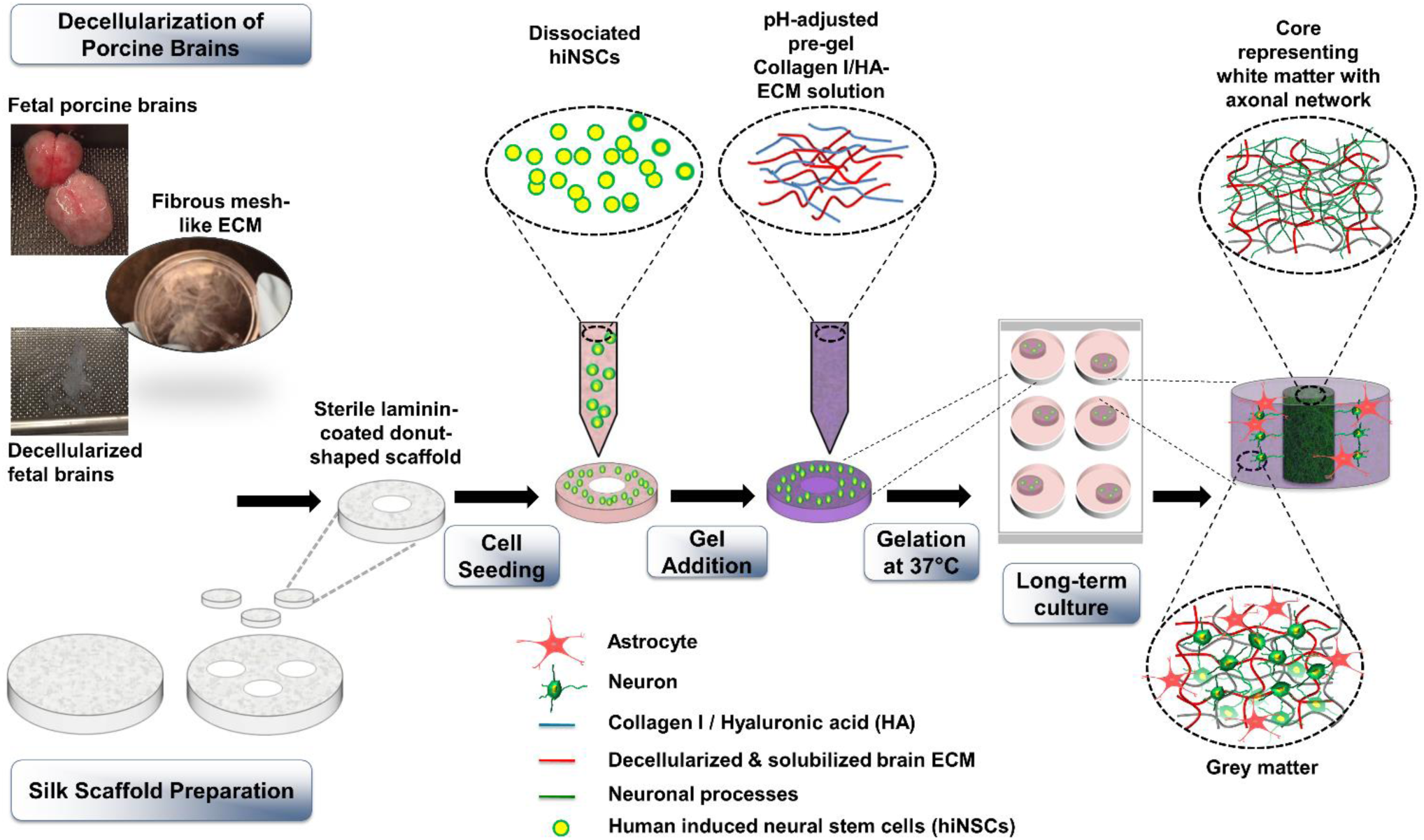
Culture of human induced neural stem cells in 3D *in vitro* bioengineered brain tissue constructs infused with decellularized brain ECM-collagen I/ hyaluronic acid (HA) hydrogel. The process starts with decellularization of porcine brains and silk scaffold preparation. Scaffolds are punched into 6 mm diameter constructs with a 2 mm diameter central hole. Laminin-coated scaffolds are seeded with dissociated human induced neural stem cells (hiNSCs). Decellularized ECM is mixed with either collagen I or HA solution and added to the scaffolds seeded with cells. The cell-seeded silk-scaffolds are flooded with media after complete gelation of ECM-collagen I or ECM-HA. The center of the construct shows a dense axonal network representing white matter, surrounded by the neuronal cell bodies and astrocytes representing the grey matter.

## Results

### Extracellular matrix and time-dependent differentiation of hiNSCs in 3D cultures

We tested whether the presence of brain-derived ECM cues accelerated the differentiation of hiNSCs into mature neurons and glia, particularly astrocytes, in the 3D bioengineered brain tissues. An increased density of axonal network from differentiating neurons was observed in fetal ECM-enriched constructs as shown by beta-III tubulin staining at 6 weeks, within both the axon rich central window (Figure 1a-b) and the scaffold portion of the constructs (Figure 1a-c). The axonal projections filled the central window significantly faster in the brain ECM-enriched constructs in comparison to unsupplemented collagen type I or HA, when collagen type I or HA were used as the hydrogels, respectively (Figure 1b, Supplementary Figure 1a). Additionally, time-dependent increased differentiation of hiNSCs into neurons and astrocytes was observed based on immunostaining (Figure 1d). The astrocyte population was more evident in 2-month cultures, closely following the differentiation of neurons. Star shaped astrocytes, suggestive of mature resting astrocytes, were only visible in the brain ECM-enriched constructs, particularly in 7-month cultures (Figure 1d, marked by white arrows). The structural integrity of the neurons and astrocytes was maintained throughout the culture duration within the fetal brain ECM-enriched 3D brain tissue constructs (Figure 1d, left column). In contrast, unhealthy neuronal morphologies, visible either as traces of disintegrated axons or as debris of clumped neuronal cell bodies, were present in the unsupplemented collagen type I constructs at all time points, and were particularly evident at 7 months (Figure 1d, right column, marked by white arrow heads).

**Figure 1:**
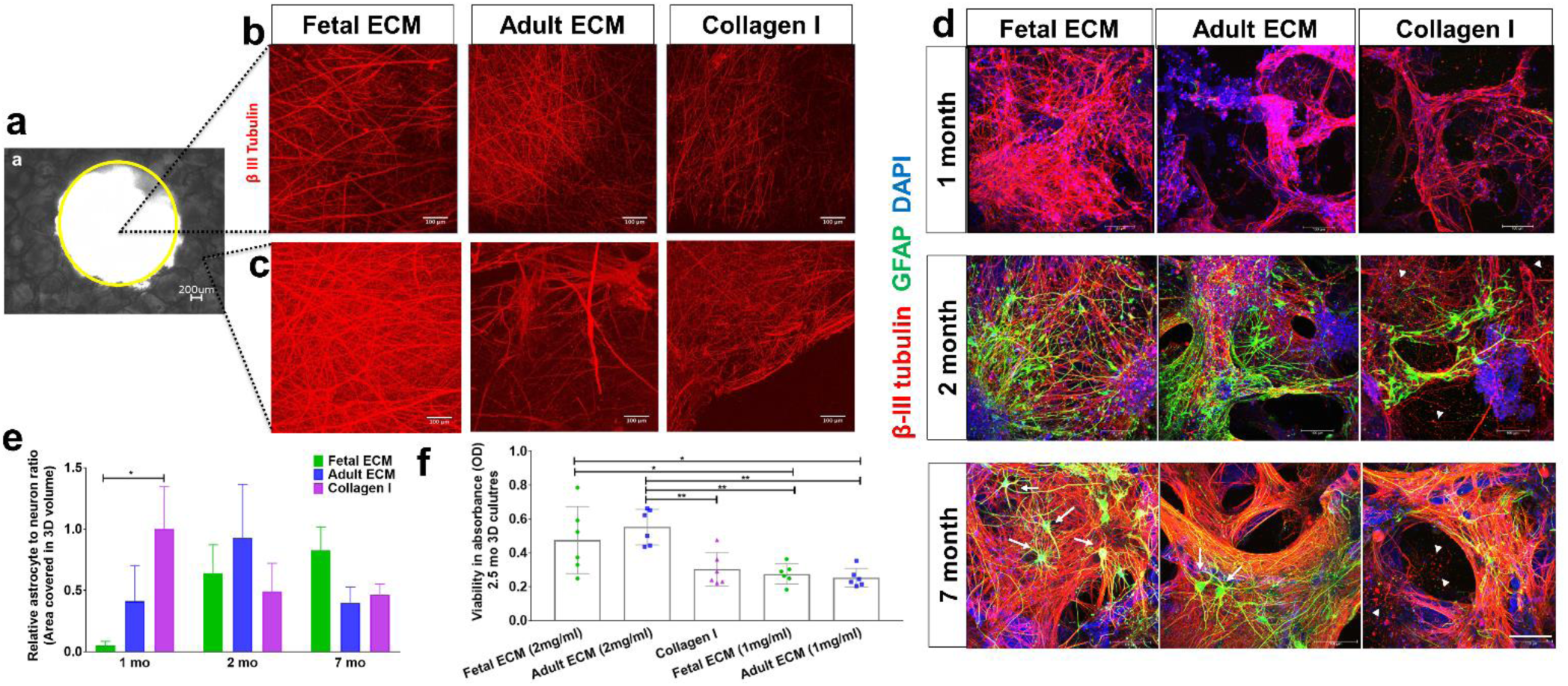
Extracellular matrix and time-dependent differentiation of human induced neural stem cells in 3D cultures. Human induced neural stem cells (hiNSCs) in silk scaffold-based 3D constructs infused with collagen I hydrogels supplemented with native porcine brain-derived ECM. (a) Brightfield image of silk scaffold with the middle circular window indicated by the yellow outline. (b) Growth of differentiating hiNSCs at 6 wk shown by β-III tubulin staining for neurons within the middle hydrogel window of the 3D donut-shaped constructs. Max projection of z-stack. Scale bar 100μm. (c) Growth of differentiating hiNSCs at 6 wk shown by β-III tubulin staining for neurons within the ring portion of the 3D donut-shaped constructs. Max projection of z-stack. Scale bar 100μm. (d) Growth and differentiation of hiNSCs at 1, 2 and 7 mo shown by β-III tubulin staining for neurons (red) and GFAP staining for astrocytes (green) across different ECM conditions. Max projection of z-stack. Scale bar 100μm. (e) Astrocyte to neuron ratio calculated by dividing the total volume in 3D confocal stacks covered by astrocytes versus neurons post image processing. Mean±SEM. One-way ANOVA with Dunnett’s post hoc test (Collagen I as control condition) at each time point on log transformed data, n=3-6. (f) Wst-1 viability assay at 2.5 mo in 3D hiNSC cultures. One-way ANOVA with Tukey’s post hoc for multiple comparisons. * p< 0.0431,** p< 0.0071.

These qualitative observations were followed by the quantification of the volume covered by neurons and astrocytes within the 3D confocal stacks (3<n<6 per condition). The volume covered by neurons was significantly greater in the fetal ECM constructs than adult ECM or unsupplemented collagen I at 1 month, while the astrocytic population was not evident (Figure 1d-e, Supplementary Figure 2). A time-dependent increase in the ratio of astrocytes to neurons was confirmed in the fetal ECM constructs with an initial surge of astrocytes at 2 months (Figure 1e). The inclusion of porcine brain-derived ECM had no toxicity as shown by viability and lactate dehydrogenase (LDH) release profiles across all conditions (Figure 1f, Supplementary Figure 3).

Targeted RNA profiling of 3D bioengineered hiNSC cultures was performed to determine the gene expression profiles of the differentiating cells (Figure 2a). Genes corresponding to neurons, ion channels/receptors involved in calcium signaling, mature resting astrocytes, toxic reactive astrocytes, trophic reactive astrocytes, and neural stem cell were assessed. In 1-month cultures of fetal brain ECM-enriched tissue constructs, there was an upregulation of mature neuronal markers including synapsin 1 (SYN1) and microtubule associated protein 2 (MAP2), and mature heathy astrocytes including excitatory amino acid transporter 1 (EAAT1), excitatory amino acid transporter 2 (EAAT2) (Sloan et al., 2017), and multiple epidermal growth factor-like domain protein 10 (MEGF10) (Chung, Allen, & Eroglu, 2015). All of these markers were higher than in unsupplemented collagen I cultures of similar age (Figure 2a). On the other hand, markers of toxic reactive astrocytes such as Serpina3 (Clarke et al., 2018) were downregulated in fetal brain ECM-enriched tissue constructs (Figure 2a). Concurrent upregulation of many different voltage gated ion channels (sodium, potassium and calcium) was evident in brain-ECM supplemented constructs, particularly with fetal brain ECM (Figure 2a), suggestive of excitability and synaptic transmission (Vacher, Mohapatra, & Trimmer, 2008).

**Figure 2:**
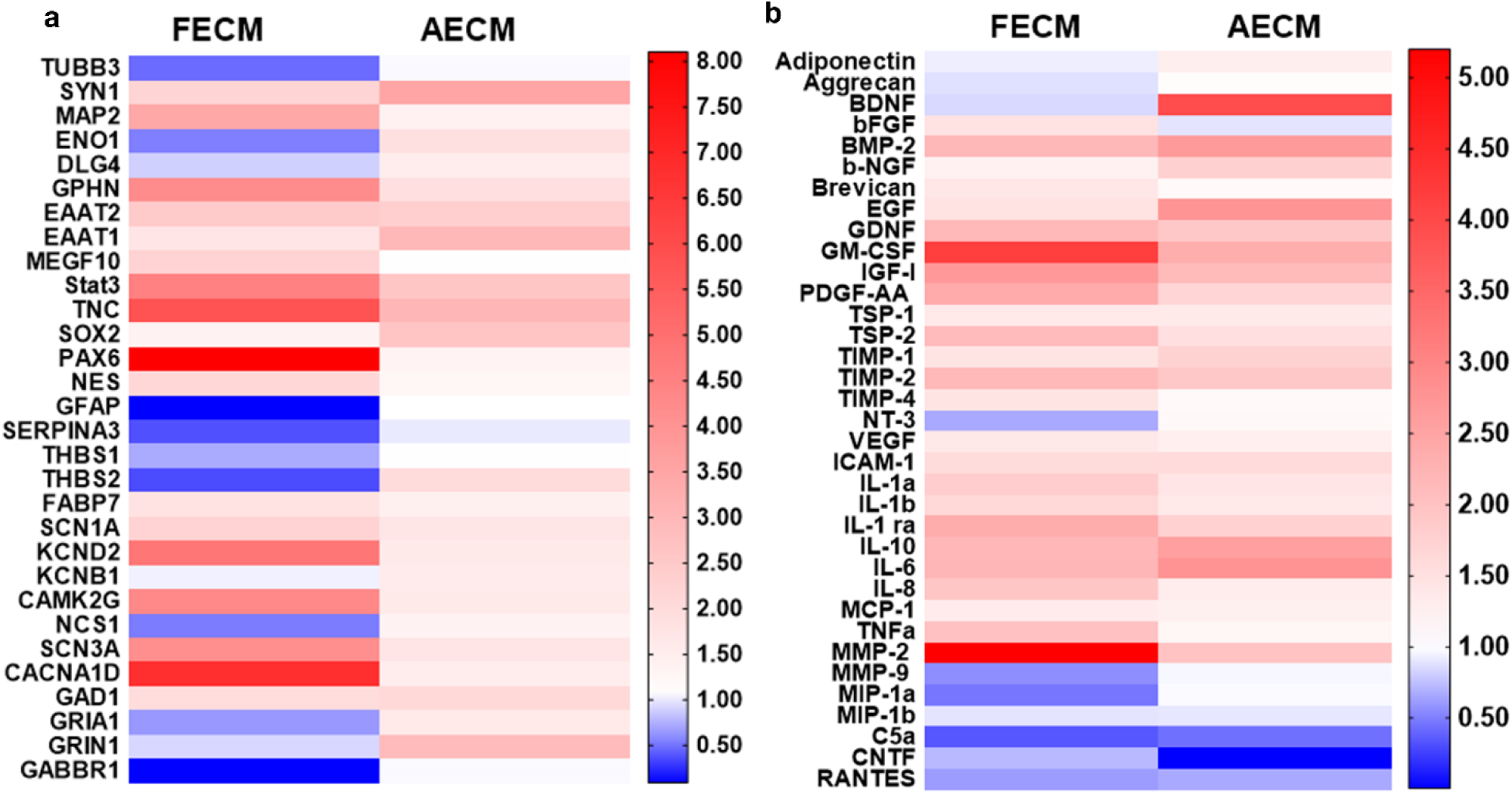
Gene expression changes in 3D bioengineered human induced neural stem cell cultures (hiNSCs) cultured in decellularized fetal or adult brain ECM. (a) Left and right panels indicate fold change in gene expression within fetal ECM and adult ECM-enriched constructs relative to collagen I condition, respectively. n=3 pooled per condition at 1 month. (b) Cytokine release profile of differentiating hiNSCs in 3D bioengineered cultures at 1 month.Media was pooled from n=7 samples per condition for the cytokine microarray. Refer to Tables 1-2 for the detailed list of genes and cytokines.

**Table 1:** List of genes tested for release in 3D differentiating human neural stem cell cultures.

|  |  |
| --- | --- |
| CACNA1D | Voltage-dependent L-type calcium channel subunit alpha-1D |
| CAMK2G | Calcium/calmodulin-dependent protein kinase type II subunit gamma |
| DLG4 | Disks large homolog 4 |
| EAAT1 | Excitatory amino acid transporter 1 |
| EAAT2 | Excitatory amino acid transporter 2 |
| ENO1 | Enolase 1 |
| FABP7 | Fatty Acid Binding Protein 7 |
| GABBR1 | Gamma-aminobutyric acid type B receptor subunit 1 |
| GAD1 | Glutamate decarboxylase 1 |
| GFAP | Glial fibrillary acidic protein |
| GPHN | Gephyrin |
| GRIA1 | Glutamate receptor 1 |
| GRIN1 | Glutamate receptor ionotropic, NMDA 1 |
| KCNB1 | Potassium voltage-gated channel subfamily B member 1 |
| KCND2 | Potassium voltage-gated channel subfamily D member 2 |
| MAP2 | Microtubule associated protein 2 |
| MEGF10 | Multiple epidermal growth factor-like domains protein 10 |
| NCS1 | Neuronal calcium sensor 1 |
| NES | Nestin |
| PAX6 | Paired box protein Pax-6 |
| SCN1A | Sodium channel protein type 1 subunit alpha |
| SCN3A | Sodium channel protein type 3 subunit alpha |
| SERPINA3 | Peptidase inhibitor Serpina3n |
| SOX2 | Transcription factor SOX-2 |
| Stat3 | Signal transducer and activator of transcription 3 |
| SYN1 | Synapsin 1 |
| THBS1 | Thrombospondin 1 |
| THBS2 | Thrombospondin 2 |
| TNC | Tenascin |
| TUBB3 | Beta-III tubulin |

**Table 2:** List of cytokines tested for release in 3D differentiating human neural stem cell cultures

|  |  |
| --- | --- |
| ACRP30 | Adiponectin |
| BDNF | Brain-derived neurotrophic factor |
| BMP2 | Bone morphogenetic protein 2 |
| b-NGF | beta- Nerve growth factor |
| bFGF | Basic fibroblast growth factor |
| C5a | Complement component 5a |
| CNTF | Ciliary neurotrophic factor |
| EGF | Epidermal growth factor |
| G-CSF | Granulocyte-colony stimulating factor |
| GDNF | Glial derived neurotrophic factor |
| GM-CSF | Granulocyte-macrophage colony-stimulating factor |
| GROa | Chemokine (C-X-C motif) ligand 1 |
| ICAM-1 | Intercellular Adhesion Molecule 1 |
| IGF-1 | Insulin growth factor-1 |
| IL-10 | Interleukin 10 |
| IL-1a | Interleukin 1 alpha |
| IL-1b | Interleukin 1 beta |
| IL-1ra | Interleukin-1 receptor antagonist |
| IL-6 | Interleukin 6 |
| IL-8 | Interleukin 8/Chemokine (C-X-C motif) ligand 8 |
| LIF | Leukemia inhibitory factor |
| MCP-1 | Chemokine ligand 2/CCL2 |
| MIP-1 beta | Macrophage Inflammatory Protein-1 |
| MIP-1a | Chemokine ligand 3/CCL3 |
| MMP-2 | Matrix metalloproteinase 2 |
| MMP-9 | Matrix metalloproteinase 9 |
| NT-3 | Neurotrophin-3 |
| PDGF-AA | Platelet growth factor |
| RANTES | Chemokine ligand 5/CCL5 |
| TGF beta 1 | Transforming growth factor beta 1 |
| TIMP-1 | Tissue inhibitor of matrix metalloproteinase 1 |
| TIMP-2 | Tissue inhibitor of matrix metalloproteinase 2 |
| TIMP-4 | Tissue inhibitor of matrix metalloproteinase 4 |
| TNFα | Tumor necrosis factor alpha |
| TSP-1 | Thrombospondin 1 |
| TSP-2 | Thrombospondin 2 |
| VEGF-A | Vascular endothelial growth factor A |

Secretome profiling was used to characterize the cytokine release profile of differentiating hiNSCs. Soluble cytokines that are known to be important in astroglial differentiation (e.g., glial derived neurotrophic factor (GDNF) (Gowing et al., 2014)) and in generating/maintaining healthy neurons/synapses (e.g., Brevican (Barros, Franco, & Muller, 2011), PDGF-AA: platelet-derived growth factor AA (Funa & Sasahara, 2014), b-NGF: beta nerve growth factor (Schuldiner et al., 2001), thrombospondins (TSPs) (Risher & Eroglu, 2012)), were released in relatively greater amounts in brain ECM-enriched constructs in comparison to unsupplemented collagen I (Figure 2b). On the other hand, many cytokines associated with reactive astrocytes, including complement component 5a (C5a), Chemokine ligand 5 (RANTES) (Choi, Lee, Lim, Satoh, & Kim, 2014) and MMP-9 (Kamat, Swarnkar, Rai, Kumar, & Tyagi, 2014), were higher in unsupplemented collagen I cultures (Figure 2b). Thus, modulation of hiNSC differentiation into neurons and astrocytes was achieved using decellularized ECM derived from specific developmental stages, with fetal brain-derived ECM resulting in overall highest upregulation and release of neuronal supporting factors.

### Chondroitin sulfate proteoglycan secretion as a marker of astrocyte maturity and reactive astrogliosis

During development, chondroitin sulfate proteoglycans (CSPGs) are transiently upregulated and produced largely by maturing neurons and astrocytes; however, during disease states reactive astrocytes exhibit sustained upregulation of CSPG expression/secretion (Siebert, Conta Steencken, & Osterhout, 2014). Relative CSPG levels can thus be utilized as an indicator of astrogliosis (Yu et al., 2012). CSPGs released in 1 week and 1-month hiNSC cultures were significantly higher in fetal brain ECM-enriched constructs compared to either HA based (Figure 3a) or pure collagen type I hydrogels (Figure 3b-c), as measured in an ELISA. There was a significantly lower level of CSPG release in the brain ECM constructs at every time point post the initial month (Figure 3c). At longer culture durations, unsupplemented collagen type I containing cultures consistently showed the highest levels of CSPG release (Figure 3c). Additionally, the known morphological changes associated with reactive astrogliosis, including rapid proliferation and overlapping of cellular regions, was primarily seen in unsupplemented collagen type I matrices (Figure 3d). Therefore, based on the CSPG release profiles and GFAP-stained cell morphology (Figure 3, Figure 1d), even the nominal presence of native brain-derived ECM supported the differentiation and maintenance of healthy astrocytes when grown long-term in 3D cultures (at least up to 7 months); as opposed to culturing in pure collagen type I matrix.

**Figure 3:**
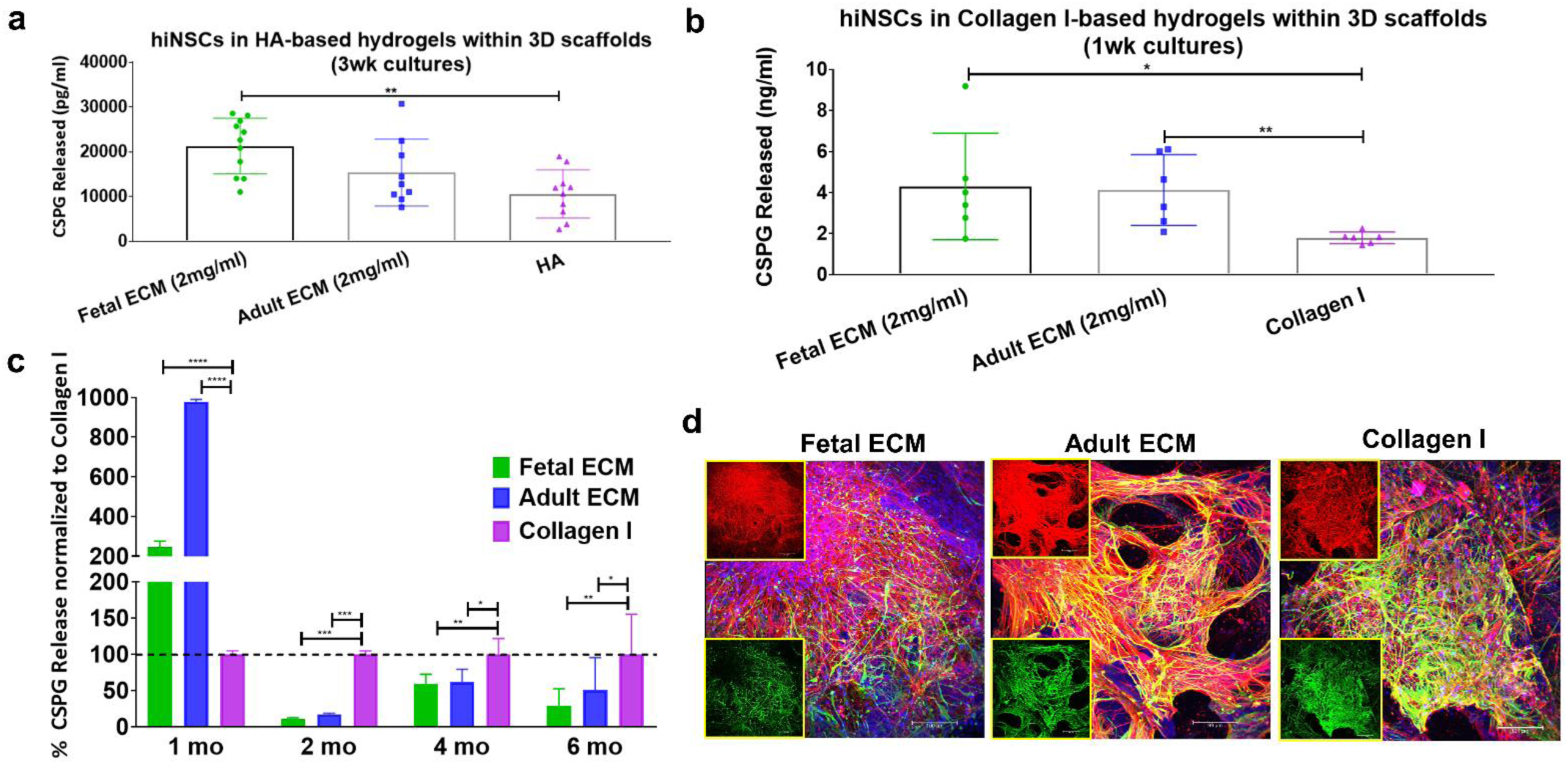
Chondroitin sulfate proteoglycans as a marker of astrocyte maturity or reactive astrogliosis. The amount of chondroitin sulfate proteoglycans (CSPGs) released in media by the cells within 3D constructs. (a) CSPGs released in media from hyaluronan-based 3D constructs at 3 wk. Unpaired two-tailed t-tests between individual pairs. (b) CSPGs released in media from collagen-based 3D constructs at 1 wk. Unpaired two-tailed t-tests between individual pairs assuming equal SD. (c) CSPGs released in media from collagen-based 3D constructs at different time points. Ordinary two-way ANOVA with Dunnett’s post hoc test and Collagen I as control condition, n=3-6. (d) Differentiating hiNSCs at 3 mo shown by β-III Tubulin staining for neurons (red) and GFAP staining for astrocytes (green) across different ECM conditions. Insets show the red and green channels separately. Max projection of z-stack. Scale bar 100μm. * p< 0.0407,** p< 0.0097, *** p< 0.0003, **** p< 0.0001.

### Extracellular matrix dependent function observed in long-term 3D cultures

Calcium wave propagation in the developing brain has been implicated in the regulation of diverse cellular differentiation, through modulation of neurotransmitter expression, and axon and dendritic morphogenesis (Rosenberg & Spitzer, 2011). Differentiated neuronal and glial populations also exhibit robust and cell-specific calcium activity, including single cell spikes or network bursts (Kapucu et al., 2012). For instance, developing neurons have been shown to have increased spontaneous burst activity during the formation of synapses and networks related to their stabilization (Hua & Smith, 2004; Kamioka, Maeda, Jimbo, Robinson, & Kawana, 1996). Thus, the spatiotemporal patterns of calcium signaling in the 3D developing cultures were assessed to decipher the role of ECM in modulating spontaneously functional networks.

Higher calcium fluorescence activity was observed in fetal ECM-enriched constructs than in adult ECM-enriched constructs and unsupplemented collagen I constructs. Specifically, greater spontaneous activity was recorded at 7 months via Fluo-4 dye (Figure 4, **Supplementary Videos 1-3**), than at 3 months (Supplementary Figure 4, **Supplementary Videos 4-6**). Moreover, the cluster analysis revealed that in fetal ECM constructs, more than 50% of the neural activity at rest was characterized by sustained oscillatory activity (cluster 1-2: Figure 4A), which was either tonic (cluster 1, Figure 4A-a) or amplitude-modulated and overlapped with spurious spikes (cluster 2, Figure 4A-b). The remaining cells, instead, remained in a quiescent state with modest oscillatory activity (cluster 3, Figure 4A-c). The percentage of clustered cells with sustained oscillatory activity patterns corresponding to clusters 1-2 was 63.3%, 55.5% (Supplementary Figure 5A-B) and 57.9% (Figure 4A), representing different replicates of fetal ECM constructs respectively. In addition, the similarity among patterns within the same cluster presented a recurrent trait (Figure 4A-d), where cells in the quiescent state (cluster 3, Figure 4A-c) or in the tonic oscillatory state (cluster 1, Figure 4A-a) had high in-cluster correlation and poor out-of-cluster correlation (cluster 1: 0.77±0.21 vs. −0.23±0.42; cluster 3: 0.84±0.16 vs. −0.09±0.41; in-cluster versus out-of-cluster, mean ± SD). On the other hand, cells showing amplitude-modulated spiking activity (cluster 2, Figure 4A) had a low in-cluster correlation value (0.34±0.43 vs. −0.39±0.35, in-cluster versus out-of-cluster, mean ± SD, Figure 4A-d). This trend was consistent across neural populations involving fetal ECM (Supplementary Figure 5A-a and 5B-a), and along with the scattered arrangement of cluster 2 cells in the construct (Supplementary Figure 6 a-c), indicated that the spontaneous spiking was decorrelated across cluster 2 cells and reflected a low-level of established connectivity.

**Figure 4:**
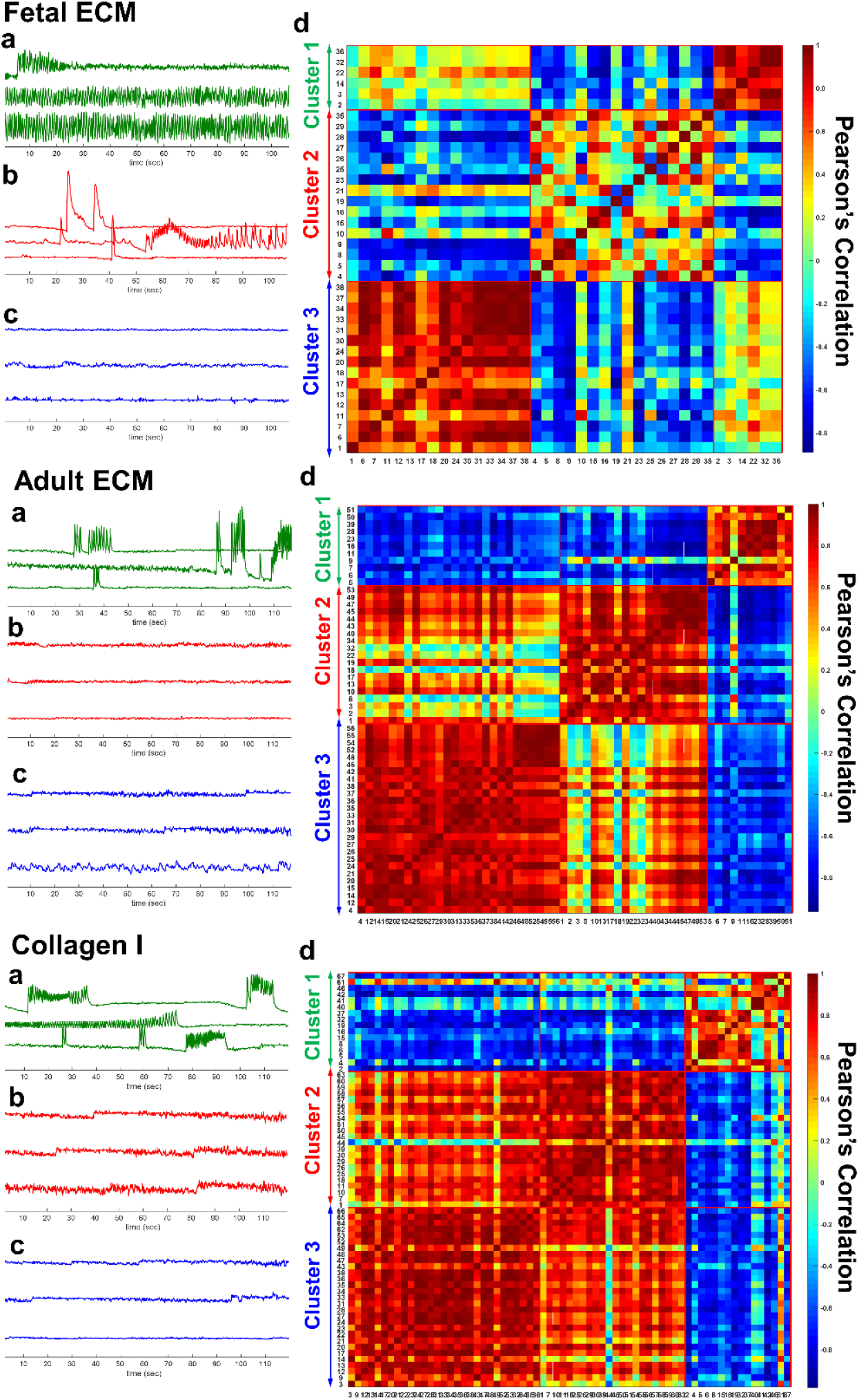
Spontaneous calcium activity in 3D cultures of differentiating human neural stem cells at 7 months. <u>Panel A:</u> cells from a construct with fetal ECM; <u>Panel B:</u> cells from a construct with adult ECM; <u>Panel C:</u> cells from a construct with unsupplemented collagen I. In each panel: (a-c) Three clusters were identified using the Louvain algorithm and three examples of ΔF/F time series per cluster are reported (a: cluster 1; b: cluster 2; c: cluster 3). (d) Pearson’s correlation coefficients between vectors *f*. There is one vector per ROI and ROIs are sorted according to the cluster position. Numbers on the edges of the matrix indicate the unique ID of the ROI.

Unlike the fetal case, constructs involving adult ECM presented a low percentage of clustered cells with spontaneous spiking activity (i.e., 19.6%, 40%, 27.27% corresponding to clusters 1 in Figure 4B-a, Supplementary 7A-b and Supplementary Figure 7B-b, respectively) and a majority of cells showed a generalized quiescent condition (Figure 4B-b and c), which rarely alternated with modest oscillatory activity (e.g., ΔF/F at the bottom of Figure 4B-c). This was further confirmed by the high inter-cluster similarity between cluster 2 and cluster 3 in Figure 4B-d (0.41±0.37, Pearson’s correlation coefficient, mean ± SD) and remained consistent across different neural populations (see Supplementary Figure 7A-B). Analogously, neural population growth in collagen-based constructs had low percentages of cells with spontaneous spiking activity (23.9% of the population depicted in Figure 4C [cluster 1], 22.7% and 29.0% of the population in Supplementary Figure 8A [cluster 3] and Supplementary Figure 8B [cluster 1], respectively), while the majority of cells presented either a quiescent state or low-intensity oscillatory patterns (Figure 4C, panel b-c) with high in-cluster correlation values (0.82±0.17 and 0.83±0.16, cluster 2 and 3, respectively, in Figure 4C-d), which indicated a low level of activity.

Overall, these results indicated that fetal ECM-based constructs supported neural populations with a higher fraction of active, spontaneously spiking neurons and on average, a more intense coordinated physiological activity.

### Transduction of hiNSCs with a neuron-specific reporter and genetically encoded calcium sensor for cell-specific tracking

To longitudinally track neuronal populations, differentiating hiNSCs were transduced with adeno-associated virus-dj (AAV-dj), a hybrid serotype with a higher transduction efficiency and infectivity *in vitro* in comparison to other wild type AAV serotypes (Katrekar, Moreno, Chen, Worlikar, & Mali, 2018). The transduction virus enabled the expression of eYFP (yellow fluorescent protein) throughout the cell volume under the synapsin promoter, such that the arising mature neuronal populations could be tracked over time. Additionally, a genetically encoded calcium sensor, jRCaMP1b, was expressed in the differentiating hiNSCs under the synapsin promoter. The red-shifted calcium sensor, jRCaMP1b, was particularly chosen for several advantages; brighter/stable long-term expression over GCaMP6, imaging capability at greater depths with reduced photodamage, greater sensitivity and dynamic range before saturation (Dana et al., 2016). This dual transduction enabled label-free tracking of mature neurons over time in the cultures, while simultaneously allowing for visualization of calcium levels (Figure 5a). We confirmed the presence of mature neurons expressing eYFP and jRCaMP1b at both 3 and 6 months in the 3D bioengineered cultures across all ECM conditions (Figure 5b upper and lower panels, respectively). Qualitatively, the neuronal networks were more intact and structurally robust in the fetal ECM-enriched constructs, with overall higher baseline calcium levels and increased network density at 6-months (Figure 5b, lower panel). Thus, differentiating neuronal populations were successfully transduced with dual viral constructs to specifically track these populations.

**Figure 5:**
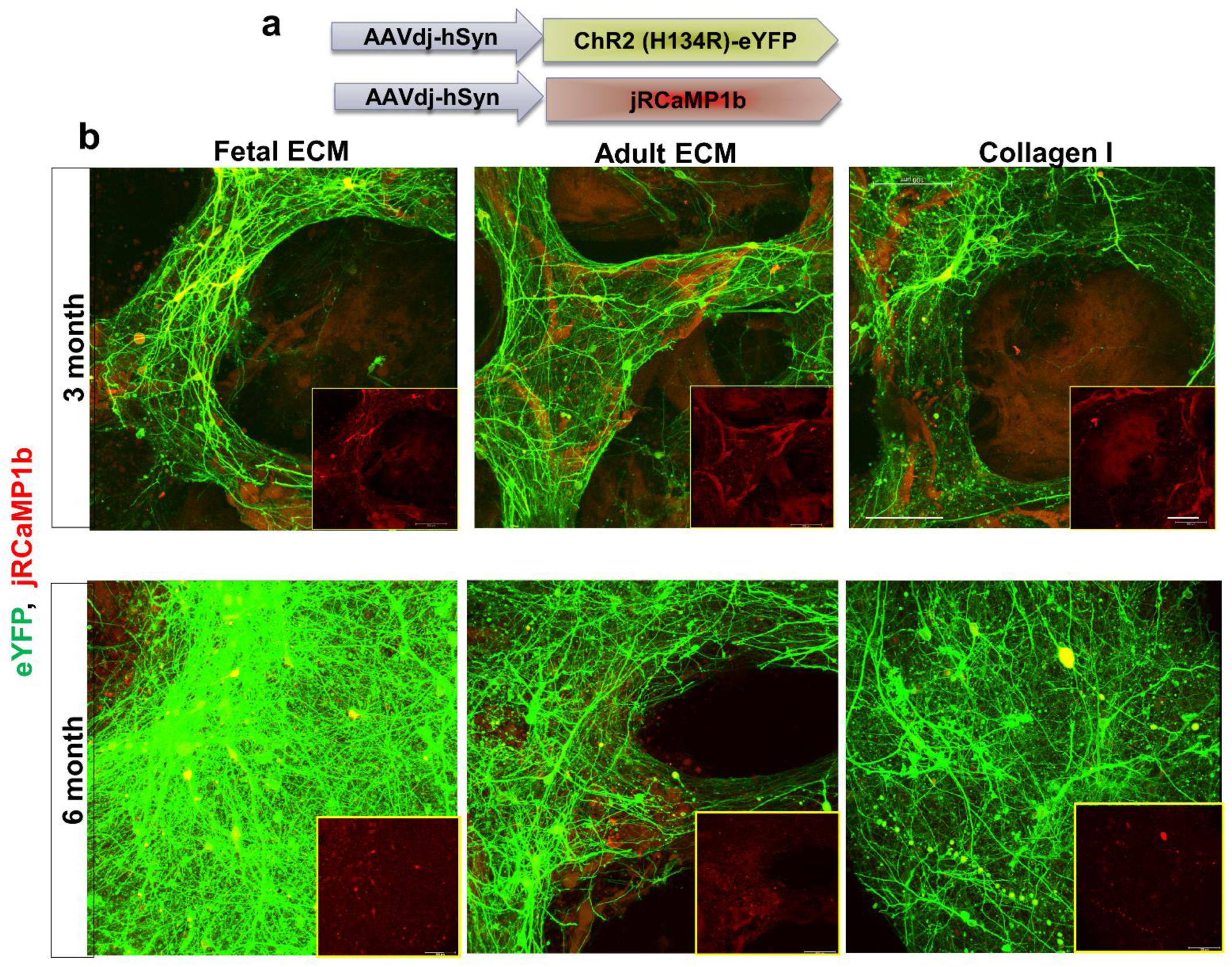
Transduction of differentiating human induced neural stem cells for tracking mature neurons. (a) High transduction efficiency virus, AAV-dj for expression of eYFP throughout the cell volume, a channelrhodopsin ChR2 (H134R) and calcium sensor jRCAMP1b under the synapsin promoter. (b) eYFP (green) and jRCAMP1b (red) expression at 3 and 6 mo in mature neurons differentiated from hiNSCs in silk scaffold-based 3D constructs infused with collagen I hydrogels supplemented with native porcine brain-derived ECM. Insets show jRCAMP1b channel only. Max projection of Z-stack. Scale bar 100μm.

### Composition analysis of decellularized brain extracellular matrix

We sought to characterize the composition of fetal and adult brain matrices, which potentially contribute to the phenotypic changes in differentiation capacity and functionality that were observed upon culture of hiNSCs in these matrices. Through Liquid Chromatography-Mass Spectrometry (LC/MS), we confirmed that a complex composition of brain ECM was maintained post-decellularization. The components varied in relative amounts between fetal versus adult ECM, but included both fibrous proteins (up to 8 types of collagens, fibronectin, laminin), glycoproteins (nidogen, fibulin, fibronectin, fibrillins) and non-fibrous proteoglycans (heparin sulfate proteoglycans/HSPGs, biglycan, mimecan, lumican) (Figure 6a). Fibrillins were higher in fetal brain-derived ECM (Figure 6b). Furthermore, biglycans were enriched in the decellularized fetal brain ECM, when tested over multiple different extractions (Supplementary Figure 9a).

**Figure 6:**
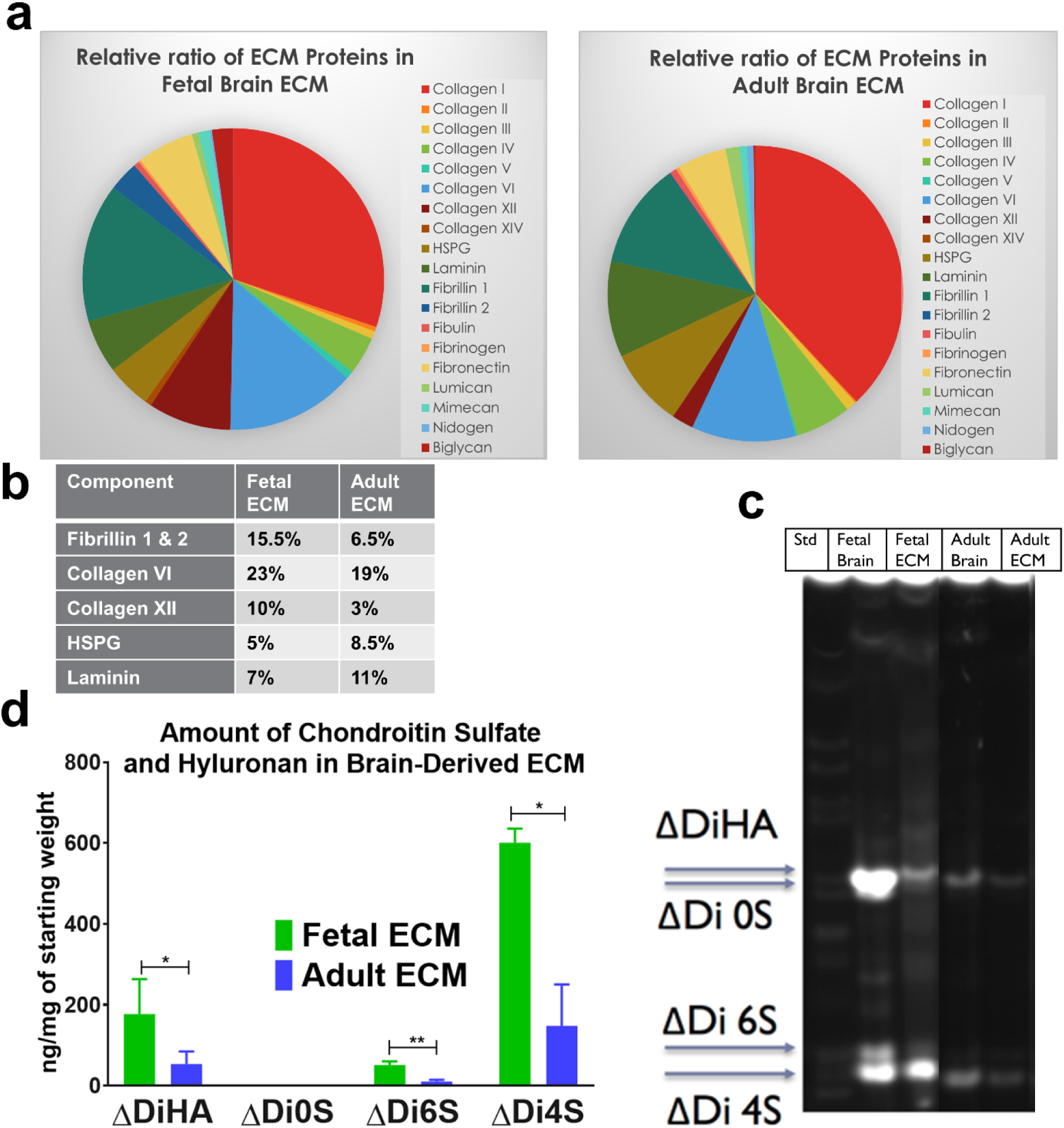
ECM composition analysis using LC/MS and fluorescence assisted carbohydrate electrophoresis. (a) Relative ratios of ECM proteins in fetal versus adult porcine brain decellularized matrix. (b) Table indicating the relative percentages of a few select ECM components in decellularized brain matrix. (c) Overall higher quantity of GAGs present in fetal brain ECM, as opposed to adult brain ECM. Unpaired two-tailed t-tests on log-transformed data between individual pairs, n=3, *p<0.0439, **p=0.0040. (d) Characterization of decellularized fetal & adult brain ECM in comparison to fetal and adult whole brains by FACE. Chondroitin sulfate (CS) and hyaluronan (HA) bands in the fetal & adult porcine brain ECM. ΔDiHA: hyaluronan, ΔDi0S: non-sulfated CS, ΔDi6S: 6-sulfated CS, ΔDi4S: 4-sulfated CS.

Considering a significant loss of GAGs during sample processing for LC/MS, we resorted to a specialized technique, Fluorescence Assisted Carbohydrate Electrophoresis (FACE), for GAG compositional analysis of the decellularized brain ECM (Calabro et al., 2001; Midura, Cali, Lauer, Calabro, & Hascall, 2018). FACE analysis of whole brain tissue indicated a greater amount of GAGs present in fetal versus adult brain (Figure 6c-d, Supplementary Figure 9b-d). This trend was maintained in the decellularized brain, with fetal ECM indicating significantly higher levels of GAGs versus adult ECM, specifically 4-sulfated chondroitin sulfate (4S-CS), 6S-CS and HA (Figure 6c-d). Additionally, HSPGs were present in higher amounts in fetal ECM (Supplementary Figure 9d).

## Discussion

There has been a growing interest in the use of *in vitro* 3D brain-like tissue systems for the study of neurological diseases using patient-derived stem cells (Lancaster et al., 2017; S. P. Pasca, 2016; Quadrato et al., 2016; Watson, Kavanagh, Allenby, & Vassey, 2017). Some of the limitations of these systems include inconsistent and slow co-differentiation of the cells towards neurons and supporting cells, such as astrocytes. Notably, Matrigel-based ECM hydrogels utilized for brain organoids, the current state of the art technology in the field, are poorly defined; this leads to variable differentiation effects such as inconsistencies in cortical layer formation and represented brain regions, including the presence of non-ectodermal identities (S. P. Pasca, 2018). Although, regionally specific brain ECM networks are reported to play major roles in generation of structural and functional diversity, little is known about how ECM is developmentally regulated in the brain (Dauth et al., 2016; S. P. Pasca, 2018). To our knowledge, the differentiation effects of fetal and adult brain ECM on human NSCs have not been characterized previously. Our previous study laid the foundation for the instructive role of decellularized fetal brain ECM in boosting primary mice neuronal culture (Sood et al., 2015). The current study investigated the composition and role of developmental stage-sourced brain ECM for enhanced and functional maturation of human induced neural stem cells (hiNSCs) into a co-culture of healthy neurons and astrocytes.

We sought to investigate the effects of developmentally sourced brain ECM in 3D bioengineered tissue systems. The advantages of a silk scaffold-based model proposed here over previously generated scaffold-free organoid cultures include structurally robust long term cultures without necrosis in the core, ease of handling and reproducibility, and segregation of gray matter and white matter with ease of monitoring neural network formation over time. We hypothesized that 3D brain constructs generated using brain-derived ECM would create a more physiologically relevant microenvironment conducive to growth and maturity of hiNSCs. We utilized hiNSCs directly reprogrammed from dermis-derived cells instead of induced pluripotent stem cells (iPSCs) because they differentiate rapidly and efficiently into both neurons and glia, without the need for lengthy protocols required for iPSC differentiation that vary in efficiency (Cairns et al., 2016). We report functional networks with enhanced maturation of neurons, predominantly in fetal brain ECM-enriched cultures at both 3 and 7 months. These functional results correlated with the upregulation of various ion channels in fetal ECM cultures.

Achieving a mature astrocyte population with temporal and physiological relevance that recapitulates the phenotype of healthy astrocytes *in vivo* is crucial. Greater differentiation of hiNSCs into healthy astrocytes was expected in the presence of native brain-derived ECM due to the neuroinductive biochemical cues, leading to the generation of a more representative 3D *in vitro* model. Indeed, we report reactive astrocyte morphology, and consistent upregulation of CSPGs in cultures lacking brain-derived ECM. This is potentially the result of reactive astrogliosis in the pure collagen I cultures at longer durations, because reactive astrocytes have been predominantly indicated for CSPG upregulation and sustained secretion in adult central nervous system (Cua et al., 2013; Pantazopoulos, Woo, Lim, Lange, & Berretta, 2010). For instance, reactive astrocyte-derived CSPG subtypes including versican V2, neurocan and phosphocan have been shown to hinder axonal growth in spinal cords of amyotropic lateral sclerosis (ALS) patients (Mizuno, Warita, Aoki, & Itoyama, 2008). In healthy cultures, CSPGs are expected to be lowered and stabilized mainly in perineuronal nets (PNNs); this is following the initial surge and in relevance to brain development where maturing neurons/astrocytes transiently upregulate CSPGs (Mueller, Davis, Sovich, Carlson, & Robinson, 2016). This was the case for brain ECM-enriched cultures, which also indicated the presence of morphologically healthy mature astrocytes (Cullen, Stabenfeldt, Simon, Tate, & LaPlaca, 2007). Additionally, we approximated the subtypes of differentiated astrocytes within brain ECM-enriched 3D bioengineered brain cultures via gene expression arrays and secretome profiling. Resting mature astrocytes identified by markers such as glutamate transporters (EAAT1, EAAT2) and phagocytic genes (MEGF10) (Sloan et al., 2017) rarely proliferate; but assume a reactive morphology and distinct markers upon activation either towards the toxic pro-inflammatory A1 or the trophic A2 subtypes (Liddelow & Barres, 2017). The existence of these two polarized populations of reactive astrocytes has been postulated, similar to macrophage polarization to subtypes M1 and M2 [56]. The pro-inflammatory A1 reactive astrocytes lose normal astrocytic functions such as neuronal outgrowth, synaptogenesis and phagocytosis, and instead contribute to neuronal death by glial scar formation and by releasing toxic factors (Clarke et al., 2018; Liddelow et al., 2017). The secretome profiling of the 2 subtypes have indicated preferentially higher release of thrombospondins from the A2 trophic astrocytes, which was noted to be the case for brain ECM containing cultures. Notably, the peptidase inhibitor serpina3a, which is postulated to be a specific marker of A1 reactive astrocytes (Lee et al., 2009; Zamanian et al., 2012), was downregulated within fetal brain ECM cultures.

Additionally, our results indicated that differentiated neurons appeared early, while astrocytes arose with increasing time in culture. The initial surge in neuronal maturation within the fetal ECM-enriched constructs leveled off at later time points. These responses are in line with the known switch towards astroglial differentiation of a multipotent cell, which initially gives rise to neuronal precursors through changes in receptor expression (Wiese et al., 2012). Moreover, maintenance of a stable astrocyte-to-neuron ratio with relevance to known *in vivo* values (∼1.4), is critical in *in vitro* brain tissue models, since this ratio is dynamic initially during development and known to increase in disease states (Nedergaard, Ransom, & Goldman, 2003). We report an astrocyte to neuron ratio with *in vivo* relevance in the generated *in vitro* brain tissue models when supplemented with fetal brain-derived ECM.

Delineating the basis for the divergence of observed cellular responses through the biochemical analysis of the brain-derived ECM may have implications for generating 3D *in vitro* brain disease models that are more representative with sufficient maturity levels. For instance, we noted overall higher amounts of fibrillins (Figure 6b) and biglycans in fetal ECM (Supplementary Figure 9a), which could present a potential mechanism for control of reactive astrogliosis and warrants further inquiry. Fibrillin 1 expression is developmentally regulated and it acts as a reservoir for transforming growth factor-β (TGF-β) in ECM. Downregulation of fibrillin 1 has been associated with increased TGF-β signaling (Burchett, Ling, & Estus, 2011). TGF-β on the other hand is a known inducer of reactive astrogliosis (Yu et al., 2012). We postulate that the fibrillins present in decellularized brain ECM could potentially harbor specific growth factors, such as TGF-β, and thus, control reactive astrogliosis in long-term 3D cultures enriched with brain ECM. Moreover, biglycan proteoglycans have been noted for their role in maintaining synaptic stability (Nastase, Young, & Schaefer, 2012). We further attribute the favorable effects of fetal brain ECM on the maturation of neural stem cells to the fact that this prenatal brain matrix allows for plasticity during development. Similarly, decellularized zebrafish brain, known for its remarkable plasticity and CNS regeneration capability, was recently shown to promote rat cortical neuronal viability and network formation in a scaffold-based culture system (S. M. Kim, Long, Tsang, & Wang, 2018).

In our 3D bioengineered cultures, collagen type I was mainly used as a base matrix into which brain ECM was incorporated, as it was deemed most stable for long term culture and best for neuronal growth through screening of different commercially available matrices (Sood et al., 2015). We also tested a custom tyramine-HRP crosslinked HA hydrogel supplemented with decellularized brain ECM (Supplementary Figure 1 b/d); however, it underwent rapid degradation. Further fractionation and analysis of ECM components will be needed to decipher the role of specific components towards differentiation of neural stem cells. The scaffold-based 3D tissue system is well suited to undertake studies for deciphering the role of specific ECM components during brain development or disease.

Furthermore, through transduction of differentiating neurons with an opsin, e-YFP reporter and genetically encoded calcium sensor (GECI), we demonstrated that this system is amenable to live tracking as well as all optical interrogation of cells over long-term cultures. Similar approaches could be used to place an opsin and GECI under a healthy astrocyte promoter, with distinct spectral profiles from those placed under the neuronal promoter synapsin; which would eventually enable studies of cell-cell interactions towards the generation of network patterns and dissect cell-specific signaling. Moreover, the resulting scaffold-based brain tissue model with concomitant presence of neurons and astrocytes is highly tunable from a structure-morphological perspective. This feature enables gray/white matter segregation and presents potential for spatially controlled introduction of morphogens or other cell types in future iterations, such that their roles in neurodegenerative diseases can be investigated.

## Conclusions

First, we report healthy mature astrocyte morphology supported by relevant gene expression/cytokine release, and downregulation of CSPGs in cultures supplemented with brain-derived ECM. Our results indicate that differentiated neurons appeared early, closely followed by astrocytes. Such systems would be useful to investigate neurodegenerative disorders, many of which have implicated reactive astrogliosis as a cause or consequence. Next, the combined functionality of cells differentiated from hiNSCs in the 3D bioengineered tissue model, revealed greater overall spontaneous activity at 7-month versus 3-month cultures. Clear differences were observed across ECM conditions, including more active clusters overall, increased coordinated activity, highest concurrent upregulation of voltage gated ion channels and downregulation of markers of toxic reactive astrocytes in the fetal ECM-enriched constructs. Furthermore, live tracking of differentiating neurons in long-term 3D cultures was achieved using genetically encoded biosensors.

This is the first study to examine the composition of decellularized brain ECM from different developmental stages, specifically glycosaminoglycans (GAGs), and to identify native stimulatory cues relevant for functional maturation of hiNSCs over long term in 3D bioengineered brain constructs supplemented with decellularized fetal and adult porcine brain ECMs. Moreover, the combination of the proposed measurements of neurons and/or astrocytes with functional optogenetic interrogation in future iterations holds the potential to help unravel cell-matrix crosstalk and to understand the interactions between cell types in healthy versus diseased states. Altogether, the knowledge gained has the potential to enable the development of brain-specific biomaterials for generating physiologically-relevant 3D *in vitro* brain models.

## Materials and Methods

### 3D Bioengineered Brain Model with hiNSCs

Assembly of the bioengineered cortical tissue was performed as previously described with further optimization (Chwalek, Sood, et al., 2015). Briefly, porous silk 3D scaffolds were coated with 0.05-0.5 mg/mL laminin (Sigma-Aldrich) either overnight at 4°C or for 2 h at 37°C. The scaffolds were incubated in media at 37°C for at least 30 mins to equilibrate the scaffolds for cell seeding. Expanding hiNSCs were lifted off mouse embryonic fibroblasts (MEFs) using TrypLE Select and centrifuged at 3,000 rpm for 1.5 mins. The cell pellet was resuspended in hiNSC expansion media consisting of KnockOut Serum Replacement DMEM (Thermo Fisher, cat#10829-018), GlutaMax (Thermo Fisher, cat#35050-061), KnockOut SR (Thermo Fisher, cat#A1099202), Antibiotic-Antimycotic (Thermo Fisher, cat#15240-062) and 2-mercapto (Thermo Fisher, cat#21985-023), bFGF Basic (Thermo Fisher, cat#PHG0024), as previously defined (Cairns et al., 2016). The resuspended cell solution was vortexed and passed through a 40 µm filter to achieve single cell suspension. The resulting single cells obtained from hiNSC colonies were seeded on the 3D ring-shaped silk scaffolds at a concentration of 0.5-1 million in 100 µl volume per scaffold. After overnight incubation of 100 µl hiNSC concentrated cell suspension per scaffold in a 96-well plate to maximize cell attachment to the laminin coated silk, the unattached cells were washed away with the hiNSC expansion media. Next, the hiNSC cell-seeded scaffolds were infused with either 3 mg/mL rat tail collagen type I (Corning), a commonly used matrix or with collagen-native brain ECM composite hydrogels.

For the generation of ECM-collagen I hydrogels, porcine brain ECM from different developmental stages were obtained via a previously developed decellularization process (Sood et al., 2015). Lyophilized ECM was solubilized with 1 mg/mL pepsin from porcine gastric mucosa (Sigma-Aldrich) in 0.1N hydrochloric acid (Sigma-Aldrich). The solubilization time for fetal and adult ECM at room temperature was approximately 16 and 24 h, respectively. Once solubilized, the ECM was mixed with hiNSC differentiation media (Neurobasal media supplemented with 1% B27, 1% glutamax and 1% anti-anti) at a 1:1 ratio and neutralized using 1 M NaOH (Sigma-Aldrich). The neutralized ECM solution was mixed with 3 mg/mL rat tail collagen type I (Corning) for a final ECM concentration of 1,000 μg/ml or 2000 μg/ml and the gelation process started using NaOH. The ECM-collagen solution was kept on ice until complete gelation was required and was used within 2 h of preparation. For hyaluronic acid (HA) hydrogels, 5.5% tyramine-substituted HA (Lifecore) was reconstituted under sterile conditions at 10 mg/ml in ultrapure water overnight at 4°C on a shaker. To obtain HA gels of ∼1 kPa bulk modulus, final optimized concentrations of HA (4 mg/ml), horseradish peroxidase (1 U/mL of gel), hydrogen peroxide (0.005% v/v) and pH adjusted 10x DMEM (1x in gel) were mixed together on ice. The remaining volume was adjusted by addition of ultrapure water. In the case of ECM-HA hydrogels, 10x DMEM (final 1x in gel) was added to solubilized ECM, which was further pH adjusted and then mixed with the rest of the components. HA hydrogels were prepared in small volumes (∼1 ml) due to their rapid gelation time and added to scaffolds immediately. After introduction within the cell-seeded scaffolds, the gelation was completed in 30 mins at 37°C, following which more media was added to each well with the constructs. The next day, each of the ECM containing cell-seeded 3D constructs was moved to a larger well of a 24-well plate with sufficient media.

### Immunostaining and Quantification of Area Covered by Neurons versus Astrocytes

At different time points (1, 2, 7, 13 months) in 3D cultures, cells were evaluated for neuronal network density and differentiation into neurons and astroglial cells with immunostaining. The samples were fixed at different time points with 4% paraformaldehyde (PFA) solution in PBS (Santa Cruz Biotechnology). Fixation time was 20-30 mins for the 3D constructs. The cells were stained with beta-III tubulin and GFAP (Sigma-Aldrich) as markers for neurons and astrocytes, respectively. Primary antibody incubations were performed at 4°C overnight, while the secondary antibody incubations were carried out at room temperature for 2 h. The volume covered in 3D stacks was measured using a custom code generated in MATLAB. Briefly, the 3D stacks corresponding to either beta-III tubulin or GFAP were binarized, such that there are only two possible values corresponding to each pixel (black or white). The volume covered was represented as the total number of positive pixels (black) divided by the total pixels (corresponding to either black or white pixels) of all the planes in the corresponding z-stack.

### Calcium Imaging and Cluster Analysis

Differentiated cell functionality was determined using live calcium imaging at 3 and 7 months in culture. Cells seeded on 3D scaffolds were immersed in extracellular solution: NaCl 140 mM, KCl 2.8 mM, CaCl_2_ 2 mM, MgCl_2_ 2 mM, HEPES 10 mM, glucose 10 mM, pH 7.4, (all reagents from Sigma-Aldrich). Fluo-4 (Life Technologies Corporation) calcium sensitive dye was mixed 1:1 with 20% Pluronic F127 (Life Technologies Corporation). Next, Fluo-4 was diluted to a final concentration 1 μM in the extracellular buffer prewarmed to 37°C. The Fluo-4 1 μM solution was applied on the scaffolds and incubated at 37°C for 1 h. Upon incubation, the constructs were washed with the extracellular buffer to remove excess dye. The constructs were imaged using the Keyence BZ-X700. The images were taken with following setup: 15 ms exposure, 50 ms frame frequency, 512 × 512 pixels, 4 × 4 binning, 1200 frames/min at 37°C. Images were processed offline using the NIH ImageJ software suite.

Regions of interest (ROIs) were extracted automatically from a series of calcium images over time, following a two-step approach. First, the variance of the brightness of each pixel through time was computed, allowing for the generation of a heatmap. The heatmap was convolved with a 2-D Gaussian kernel with a standard deviation computed from the resolution and magnification of the images, to ensure continuity and to reduce noise. Next, the local maxima of the filtered heatmap were found and used as seed points to isolate discrete regions, each representing an ROI. Finally, fluorescence intensity time traces were plotted on the center of mass of the discrete regions (single pixel data).

For each ROI, the relative change in fluorescence was calculated as

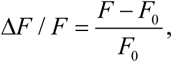

where *F* is the fluorescence time series and *F*_0_ is the baseline of *F* estimated from a ± 15 s sliding window. Due to the sparse activity, *F_0_* was calculated as the average of all values below the 80^th^ percentile in *F.* Finally, a second order Savitzky-Golay filter was applied to the Δ*F*/*F* signal to remove noise while preserving the signal frequency span.

The Δ*F*/*F* signals collected from the isolated ROIs were used for cluster analysis as in (Tang-Schomer, Jackvony, & Santaniello, 2018). Briefly, each Δ*F*/*F* was processed to compute the following features, which collectively provide a time-frequency characterization that is unique to the Δ*F*/*F* signal:

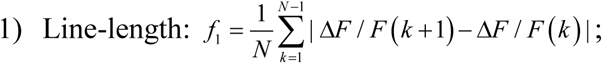

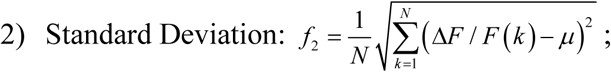

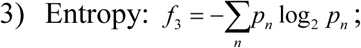

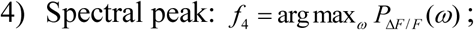

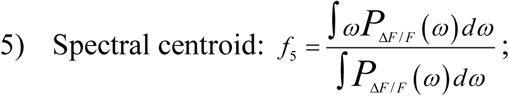

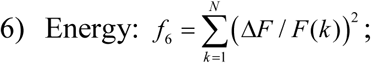

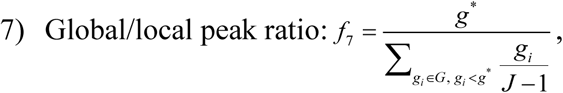

where Δ*F*/*F*(*k*) is the *k*-th sample in the Δ*F*/*F* time series, *N* is the total number of samples in Δ*F*/*F*, *μ* is the average value of Δ*F/F*, and *P*_Δ_*_F_*_/*F*_(*ω*) is the power spectrum density of the Δ*F*/*F* signal at frequency 0≤*ω*≤*F_s_*/2, where *F_s_* is the number of frames per second. To estimate the entropy (3), the sample probability function of the Δ*F*/*F* intensity values is computed and the correspondent sample probability values *p_n_* are used. The spectral peak (4), instead, is the frequency ω of the maximum power spectrum density value P_Δ_*_F_*_/*F*_(*ω*). Finally, the peak ratio (7) is estimated by computing all the local maxima (i.e., peaks) *G* = [*g_1_*, *g_2_*, *g_3_*, …, *g_J_*] of the Δ*F*/*F* signal and the absolute maximum (i.e., global peak) *g*\* among the peaks in *G*. In addition to features (1)-(7), the entropy of the squared-normalized Teager Energy vector was computed. Briefly, the Teager Energy series was computed:

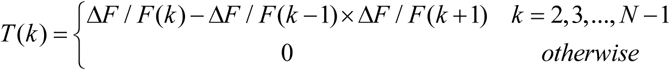

and the sample entropy is computed as:

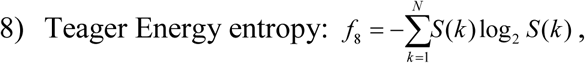

where *S*(*k*) is the squared and normalized version of *T*(*k*), i.e., 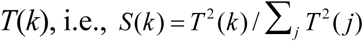. Features (1)-(8) were introduced in (Blanco et al., 2010) and provide high accuracy in distinguishing activity patterns that involve different oscillations, bursting modes, or spurious spiking.

The 8×1 vectors *f* = [*f*_1_, *f*_2_, *f*_3_, …, *f*_8_] (one vector per ROI) were finally used to cluster the ROIs, i.e., the Pearson’s correlation coefficient was computed for every pair of detected ROIs and two ROIs in one preparation were assigned to the same cluster if they had highly-correlated feature vectors. The number of clusters and the cluster assignment were determined in an unsupervised manner using the Louvain algorithm (Blondel, Guillaume, Lambiotte, & Lefebvre, 2008). The Louvain algorithm assigns the ROIs to clusters by maximizing a quantitative index that weights both the average similarity between feature vectors inside clusters and the average similarity between vectors across clusters. Because the Louvain algorithm is a locally greedy optimization algorithm, the procedure was repeated for a total of 100 optimizations and a consensus partition method was implemented as in (Lancichinetti & Fortunato, 2012) to obtain a consistent cluster partition for each culture.

### Transduction of hiNSCs with viral constructs

The starting viral titer of AAVdj-hSyn-jRCaMP1b and AAVdj-hSyn-hChR2(H134R)-eYFP (Stanford University Virus Core) was 1.78×10^13^ and 3.62×10^13^, respectively. The virus was diluted at 1:1000 dilution in cell culture media for differentiating hiNSCs. Each cell-seeded scaffold was incubated in 400 µl media containing virus and exposure to the virus was maximized by adding additional media in the same well for an entire week. One-week post-infection, virus-containing media was replaced with fresh media. Virally transduced tissue constructs were imaged with a multiphoton confocal microscope (TCS SP8, Leica) equipped with a Ti-Sapphire laser. During imaging, each sample was placed in a well of a glass bottomed (No. 1.5 coverslip) 24-well plate. The imaging chamber was maintained at 37°C and humidified along with a continuous supply of 5% CO_2_.

### Analysis of brain-derived ECM

#### Liquid Chromatography Tandem Mass Spectrometry

Lyophilized ECM samples were weighed and solubilized at 5 mg/ml in 0.1% sodium dodecyl sulfate (SDS) in PBS containing 5M urea, 2M thiourea and 50 mM Dithiothreitol (DTT). The samples were solubilized for ∼24 hrs at 4°C with gentle stirring. Following this step, the solubilized ECM samples were acetone precipitated for 2 hrs at −20 °C (Williams, Quinn, Georgakoudi, & Black, 2014). The obtained pellets following removal of the supernatant were sent for liquid chromatography tandem mass spectrometry (LC-MS/MS) at the Beth Israel Deaconess Medical Center Mass Spectrometry Core facility. The resulting spectral count data was scanned to find the most abundant ECM proteins (n=2 per ECM condition was analyzed).

#### Fluorophore-assisted Carbohydrate Electrophoresis (FACE)

Fluorophore-assisted carbohydrate electrophoresis was performed on lyophilized adult and fetal porcine brain ECM samples, using a previously developed protocol for glycosaminoglycan (GAG) analysis (Calabro, Benavides, Tammi, Hascall, & Midura, 2000). Briefly, the dry weight of the ECM samples was measured, followed by digestion in 1 mg/ml proteinase K at 60°C for 2 h. Hyaluronan and chondroitin sulfate present within the samples were digested using the following enzymes: chondroitinase ABC (Seikagaku 25mU/ml) and hyaluronidase SD (Seikagaku 2.5mU/ml). Following ethanol precipitation and proteinase K heat inactivation, the ECM samples within the resulting pellet were incubated in the enzymes at 37°C overnight. A second ethanol precipitation step was performed, and the resulting hyaluronan/CS glycans were retained within the supernatants after centrifugation. The glycans were labeled with the fluorophore 2-amino-acridone (AMAC) by incubation at 37°C for 18 h. Finally, the labeled samples and standard disaccharides were loaded onto gels at lower volumes of 2-5 µl as opposed to ∼30 µl in protein gels, to aid in band resolving. Electrophoresis was performed at a constant 500 V for 1 h 15 mins and FACE imaging was accomplished using a UVP Chemi-DocIt2 515 integrated system. Band quantifications were performed in ImageJ. To calculate the concentration of disaccharides in ECM samples, sample band intensity was divided by the intensity of the respective known standard. Relative intensities of disaccharide bands across samples were normalized by the starting dry weight.

### Statistics

Statistical analysis was performed using GraphPad Prism 7 software (GraphPad, CA, USA). All data are expressed as mean ± SD with sample sizes of *n*≥3, unless stated otherwise. The analysis methods utilized included ordinary two-way and one-way ANOVA, followed by Tukey’s *post hoc* or Dunnet’s *post hoc* test when assigning unsupplemented hydrogels as the control condition to determine the statistically significant differences for multiple comparisons, and unpaired two-tailed t-test for comparison of two groups, unless stated otherwise.

Samples were chosen randomly for the different experiments to account for any potential unavoidable biological variability. This included randomization post cell seeding within the scaffold and before allocation into different groups for ECM hydrogel addition. For clustering analysis of calcium imaging the investigator was blinded to the different groups.

## Acknowledgements

This work was funded by the US National Institutes of Health (NIH) P41 Tissue Engineering Resource Center Grant (EB002520), NIH R01 (NS092847) and NIH Research Infrastructure grant NIH S10 OD021624. Additionally, we would like to acknowledge PEG funding support for FACE training (NIH/NHLBI 1P01HL107147 Program of Excellence in Glycosciences; VC Hascall, PD/PI). We thank Mattia Bonzanni, Min Tang-Schomer, Annie Golding, Kelly Sullivan, Breanna Duffy, Whitney Stoppel and Jonathan Grasman for helpful discussions and protocols. We also thank Yu-Ting Dingle for timely help in feeding the long-term cultures and Breanna Duffy for running LC/MS samples. Additionally, we would like to extend a special thanks to Valbona Cali, Dr. Ronald Midura, Dr. Vince Hascall and Dr. Suneel Apte for training in FACE analysis at Lerner Research Institute and for their expert feedback.

## Author Contributions

D.S. and D.L.K. conceptualized ad designed the experiments, D.S. conducted the experiments, performed data analysis, data interpretation and compiled the manuscript. D.C. generated and expanded hiNSCs, advised during different stages of experimental planning, and contributed to manuscript writing. J.D. performed confocal imaging analysis, calcium imaging ROI extraction and assisted with experiments. C.R. prepared the viral constructs for neuronal transduction. C.R. and K.D. provided training at Stanford University and advised on the viral transduction protocols. L.D.B. helped conceptualize the use of developmental stage decellularized brain ECM and advised on the results. S.S. conducted cluster analysis of calcium imaging data, and contributed to clustering data interpretation and manuscript writing. D.L.K. supervised experiments and enabled all stages of manuscript preparation. All authors provided their feedback on the final manuscript.

## Data Availability Statement

All data is included within the manuscript and supplementary sections. A master source data file has been provided for Figures 1-3, 6 and Supplementary Figures 2 and 9. Source data and Matlab codes used to generate Figure 4 and Supplementary Figures 5, 7 and 8 are provided in a separate zip folder labeled-Calcium Imaging Analysis Files.

## Supplementary Methods

### Biochemical Assays

#### Viability Assay

Cell proliferation reagent WST-1 assay (Sigma-Aldrich) was performed at end time points before freezing the samples for PCR, according to the instructions provided by the manufacturer, to assess overall cell viability across different ECM conditions. Briefly, the samples were incubated for 1 h with WST-1 reagent diluted 1:10 (v:v) in culture medium, followed by a reading of the medium absorbance with plate reader (Molecular Devices) at 450 and 600 nm as the reference wavelength. Fresh medium was used as a baseline control and its average absorbance was subtracted from the value of the samples.

#### Lactate Dehydrogenase Assay

Lactate dehydrogenase (LDH) enzyme released into media by the ruptured cells, was used as a measure of cell death at different time points during the 3D culture without having to sacrifice the samples. LDH assay was performed according to the manufacturer instructions (Sigma-Aldrich). Briefly, culture medium was mixed with the assay reagents in a 1:2 ratio. Following 30 mins incubation at room temperature, the reaction was stopped by addition of 1N HCl. The absorbance readings were measured at 490 nm and 690 nm as the reference wavelengths. Fresh medium without any construct was used as a baseline control and its average was subtracted from the sample values.

#### CSPG Release ELISA

CSPGs released by the differentiating hiNSCs in media were measured using an ELISA based assay. Media samples from the 3D constructs were incubated overnight at 4°C in a 96-well immuno plate (Thermo Fischer Scientific). Alongside the sample media incubation, chicken extracellular CSPGs (Millipore) were used over a range of serial dilutions for the generation of standard curves. Following washes with PBS-tween, monoclonal anti-chondroitin sulfate antibody produced in mouse/clone CS-56, ascites fluid (Sigma) was added for overnight incubation at 4°C. After the next round of washes, HRP conjugated goat anti-mouse secondary antibody (Abcam) was incubated at room temperature for 2 h. TMB (3,3’,5,5’-tetramethylbenzidine) 1-C Substrate (Fisher Scientific) was introduced following the last round of washes with PBS-Tween. Finally, after the color developed for 10 mins at room temperature, the reaction was stopped with 1N HCl. The absorbance readings were measured at 450 nm wavelength and the fresh media readings were subtracted from the sample readings. The standard curve was utilized for calculating the quantities of CSPGs released in the different conditions and reported in pg/ml.

### qRT-PCR

Samples were flash frozen in liquid nitrogen and stored in −80°C in individual Eppendorf tubes until RNA extraction was performed. All samples were placed on dry ice during extraction, sequentially disrupted using a liquid nitrogen chilled bio-pulverizer. Between each sample, the pulverizer was wiped with 70% ethanol to remove remnants of the previous sample, and between each sample set (different tumor types), all the tools were cleaned with RNAzap. Lysis buffer was immediately added to the powdered frozen sample and placed on ice. Once all the samples were in lysis buffer on ice, a 22 gauge needle and syringe was used for sample homogenization one by one using a fresh needle and syringe every time. All the samples were spun down to remove undigested material (mainly silk scaffold) and the supernatant was transferred to clean RNAse free-Eppendorfs. Following this, the SurePrep All Prep kit (Fisher Scientific) protocol was followed until RNA was eluted from the columns. RNA concentrations were measured using nanodrop 2000 (Thermo Fisher Scientific). RT2 First Strand Kit with an incorporated gDNA removal step with buffer GE (Qiagen) was utilized for cDNA synthesis from the eluted RNA. cDNA samples were mixed with RT^2^ SYBR Green Fluor qPCR Mastermix and added to the Qiagen Custom RT2 PCR Array (including a housekeeping gene, genomic DNA control and RT control). PCR was run on BioRad CFX96. Table 1 lists the genes that were tested.

### Cytokine Arrays

Multiplex Quantibody cytokine arrays (RayBiotech) were used to semi-quantitatively compare cytokines (Table 2) released by differentiating hiNSCs cultured in the 3D bioengineered brain model with different ECM components. Small volumes of control media samples/cell culture supernatants (50 µl) from the 3D constructs were incubated in the capture antibody spotted glass slides or the membranes, along with the standards provided that corresponded to known concentrations of the targets for the Quantibody arrays. This overnight incubation was followed by another overnight step at 4°C, involving the biotinylated detection antibody cocktail. Next, streptavidin-conjugated fluorophore or HRP-streptavidin was added for 1 h at room temperature. Finally, the slide was disassembled from the removable gasket, dried and scanned using a fluorescence microarray laser scanner (Ray Biotech). Protein expression profiles of the differentiating hiNSCs across the varying ECM conditions were quantified using the Q-analyzer software (Ray Biotech). Table 2 lists the cytokines that were tested.

## Supplementary Figures

**Supplementary Figure 1:**
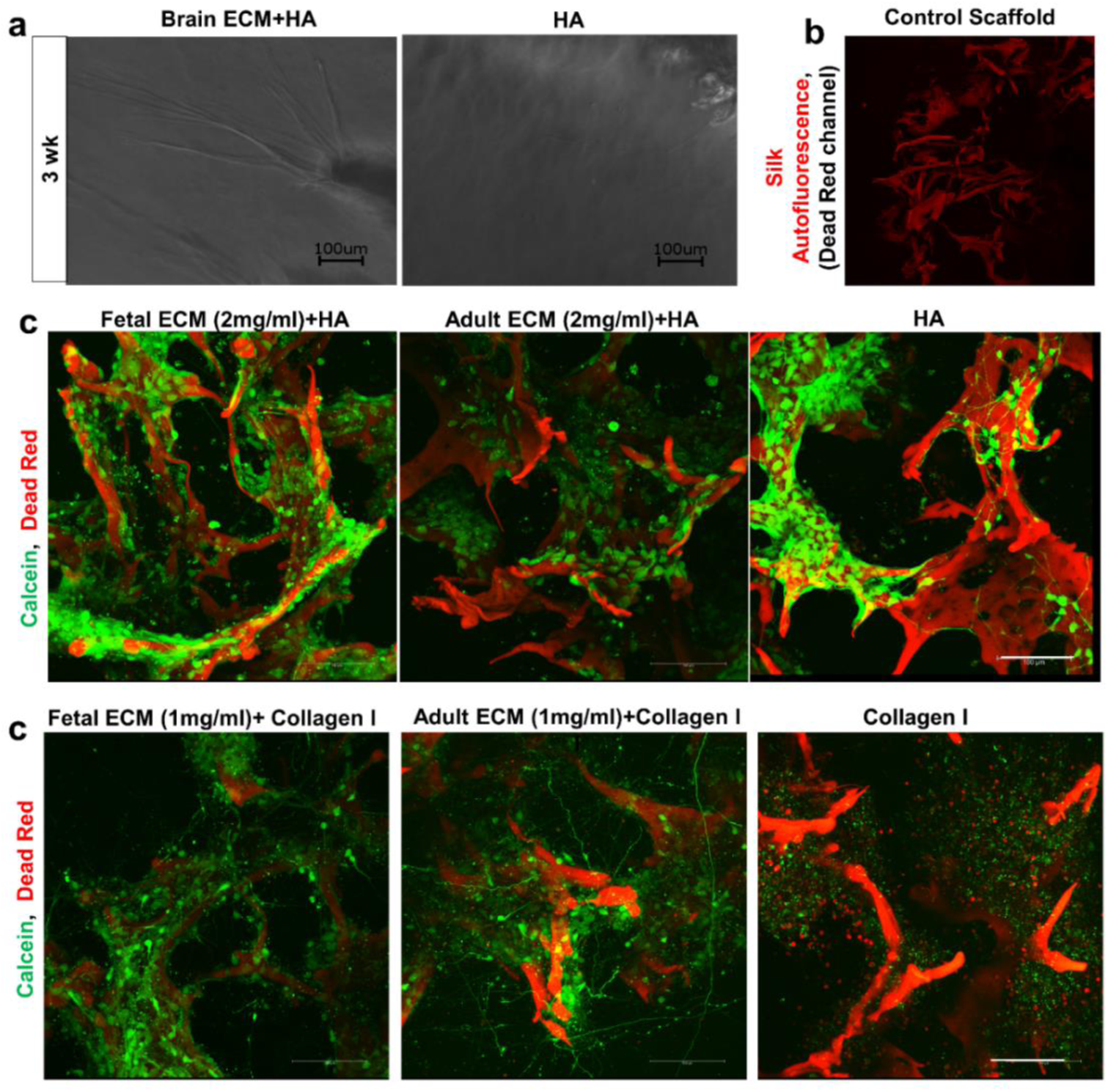
Collagen I versus hyaluronic acid (HA) based hydrogels for the 3D silk scaffold constructs. (a) Brightfield images at 3 wk indicating faster ingrowth of axons from differentiating hiNSCs towards the middle hydrogel window of the donut-shaped scaffold, when the HA hydrogel is supplemented with decellularized brain ECM. (b) Silk scaffold without cells to indicate silk autofluorescence in red channel corresponding to Dead Red. (c) Growth of differentiating hiNSCs at 10 days shown by live calcein/dead red staining within the ring portion of the 3D donut-shaped constructs infused with HA based hydrogels supplemented with native porcine brain-derived ECM in comparison to unsupplemented HA hydrogels. Max projection of z-stack, scale bar 100μm. (d) Growth of differentiating hiNSCs at 10 days shown by live calcein/dead red staining within the ring portion of the 3D donut-shaped constructs infused with collagen I based hydrogels supplemented with native porcine brain-derived ECM in comparison to unsupplemented collagen I hydrogels. Max projection of z-stack, scale bar 100μm.

**Supplementary Figure 2:**
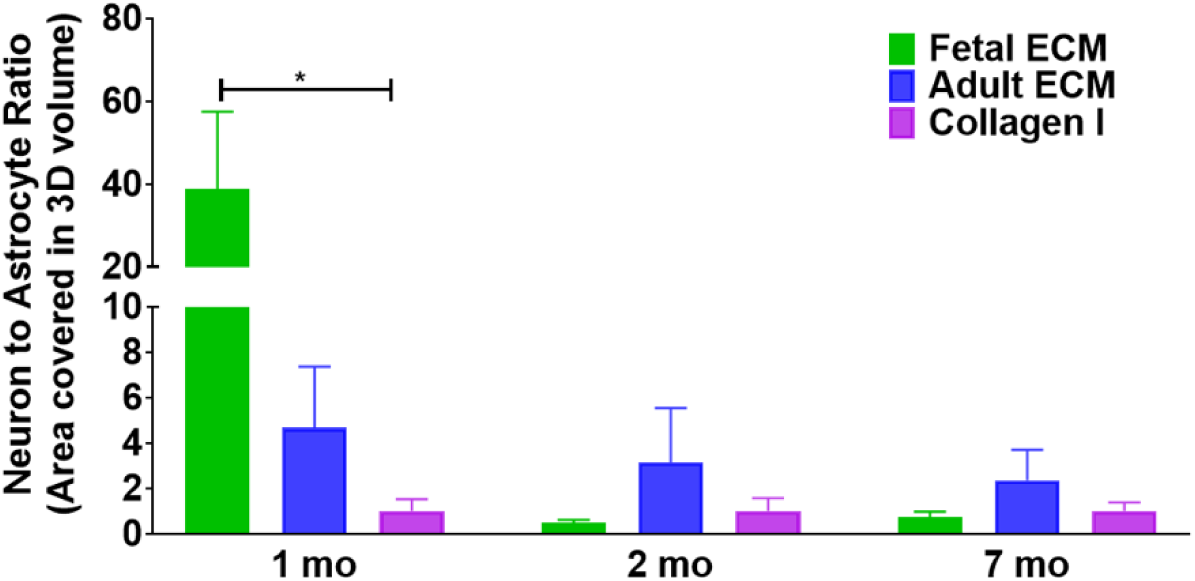
Neuron to astrocyte ratio calculated by dividing the total volume in 3D confocal stacks covered by neurons versus astrocytes post basic image processing. One way ANOVA on log transformed data with Dunnett’s posthoc at each time point, p= 0.0344, n=3-6.

**Supplementary Figure 3:**
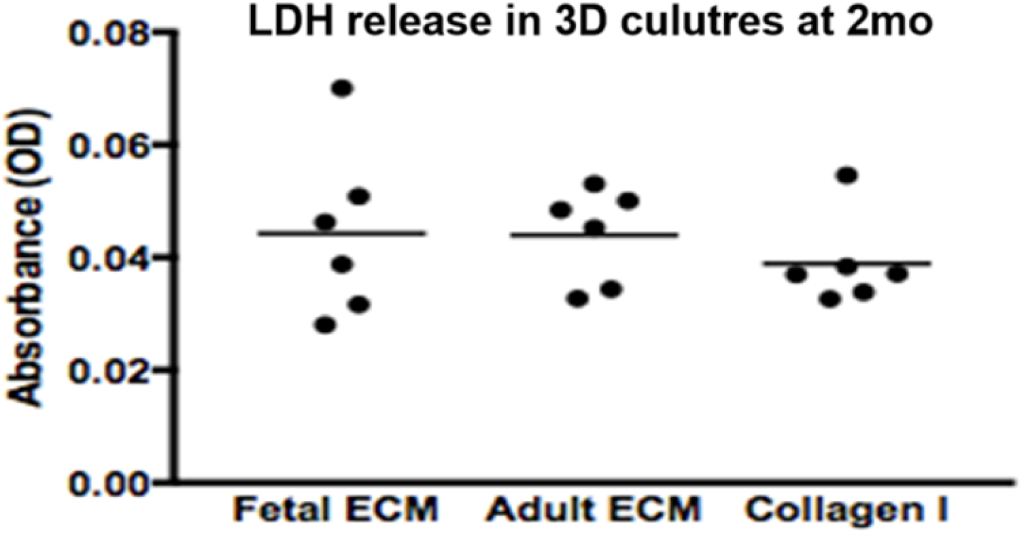
Toxicity levels at 2 mo time point in hiNSC 3D cultures as measured by LDH release in media. No statistical difference (One-way ANOVA) in comparison to collagen I when the cells were grown in the presence of decellularized ECM.

**Supplementary Figure 4:**
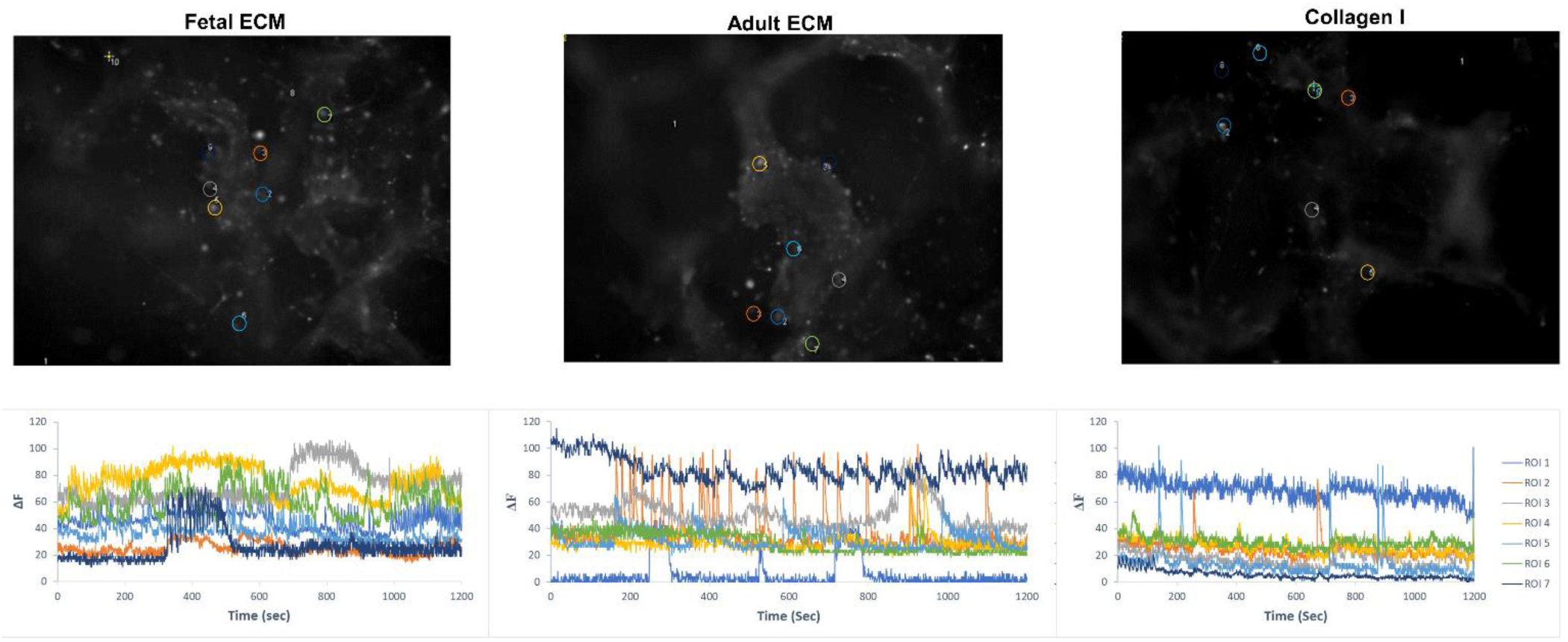
Calcium imaging of differentiating human neural stem cells in 3D cultures at 3 months. A few representative traces of spontaneous calcium activity are plotted corresponding to the regions indicated by circles in the time max projected image. ROIs were selected manually in Image J.

**Supplementary Figure 5:**
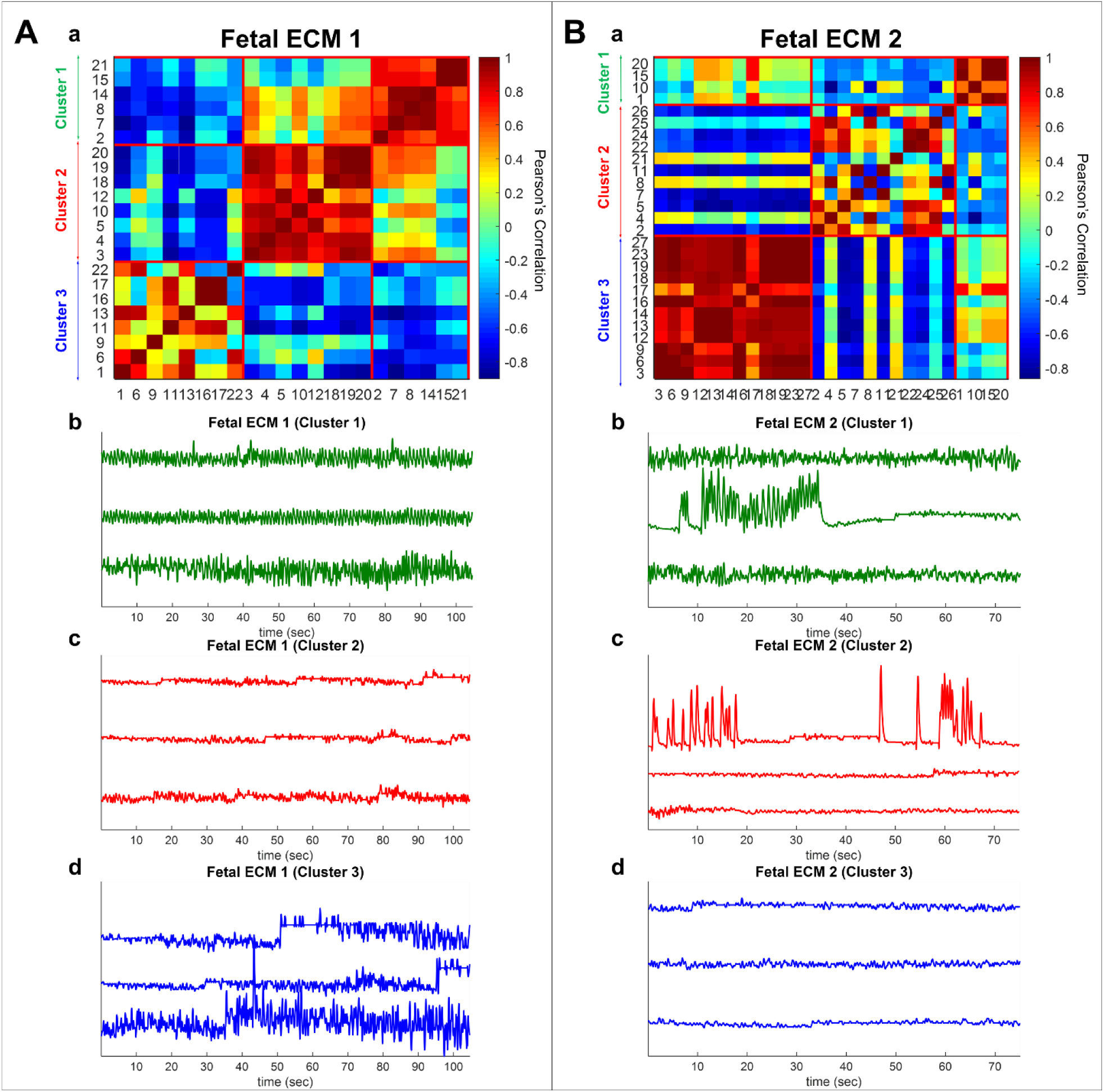
Spontaneous calcium activity in 3D cultures of differentiating human neural stem cells at 7 months; cells from fetal ECM containing constructs. Tonic oscillatory activity is captured in Cluster 1 (both panel A & B); sporadic spiking activity is captured in Cluster 3 (panel A) and Cluster 2 (panel B); quiescent state is captured in Cluster 2 (panel A) and Cluster 3 (panel B).

**Supplementary Figure 6:**
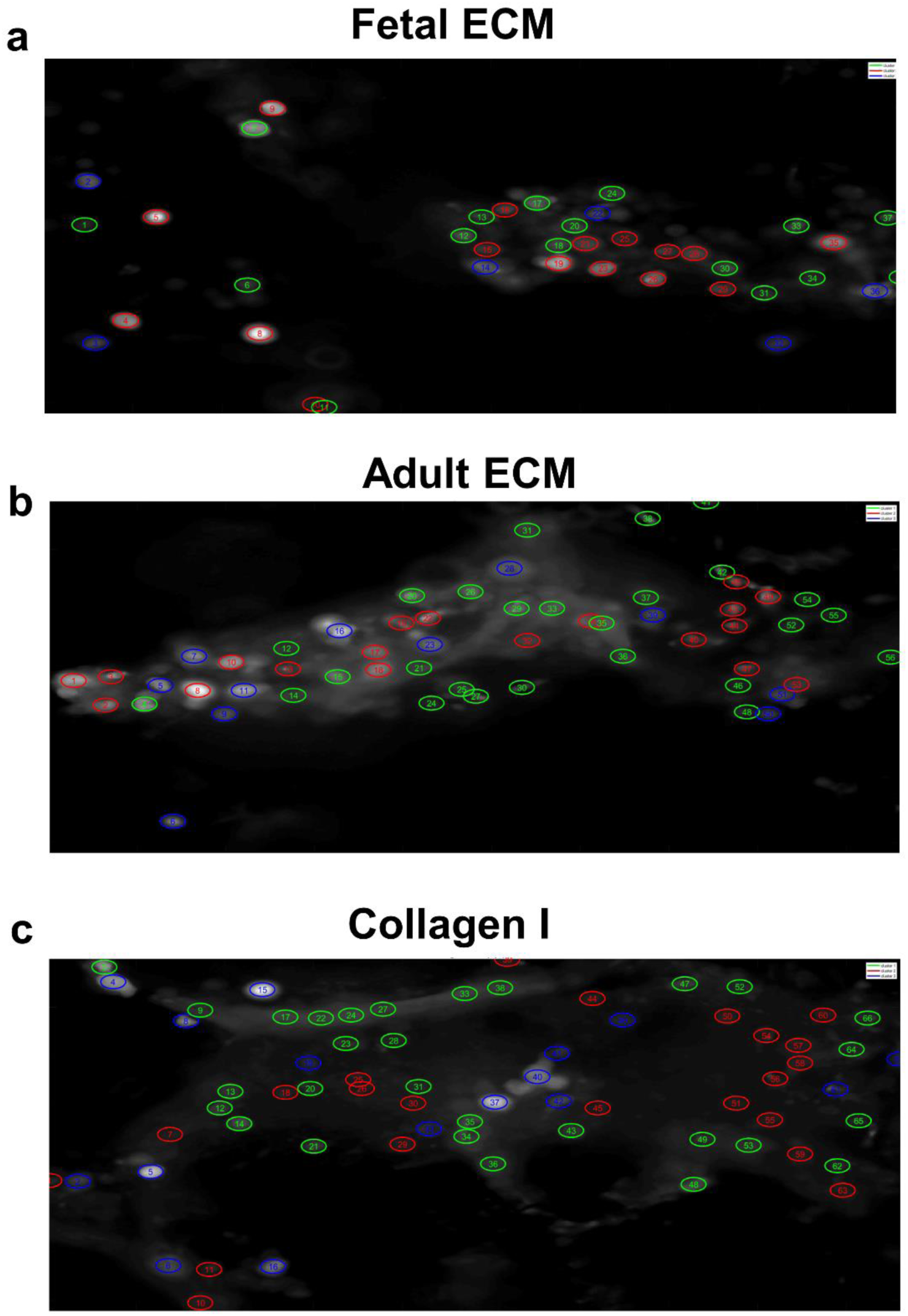
Spatial location of the ROIs in a 2D frame. Each ROI is circled and assigned with a unique ID. The color of the circles match the cluster color in Figure 4A (a-d) and the ID values are as reported in Figure 4A (d).

**Supplementary Figure 7:**
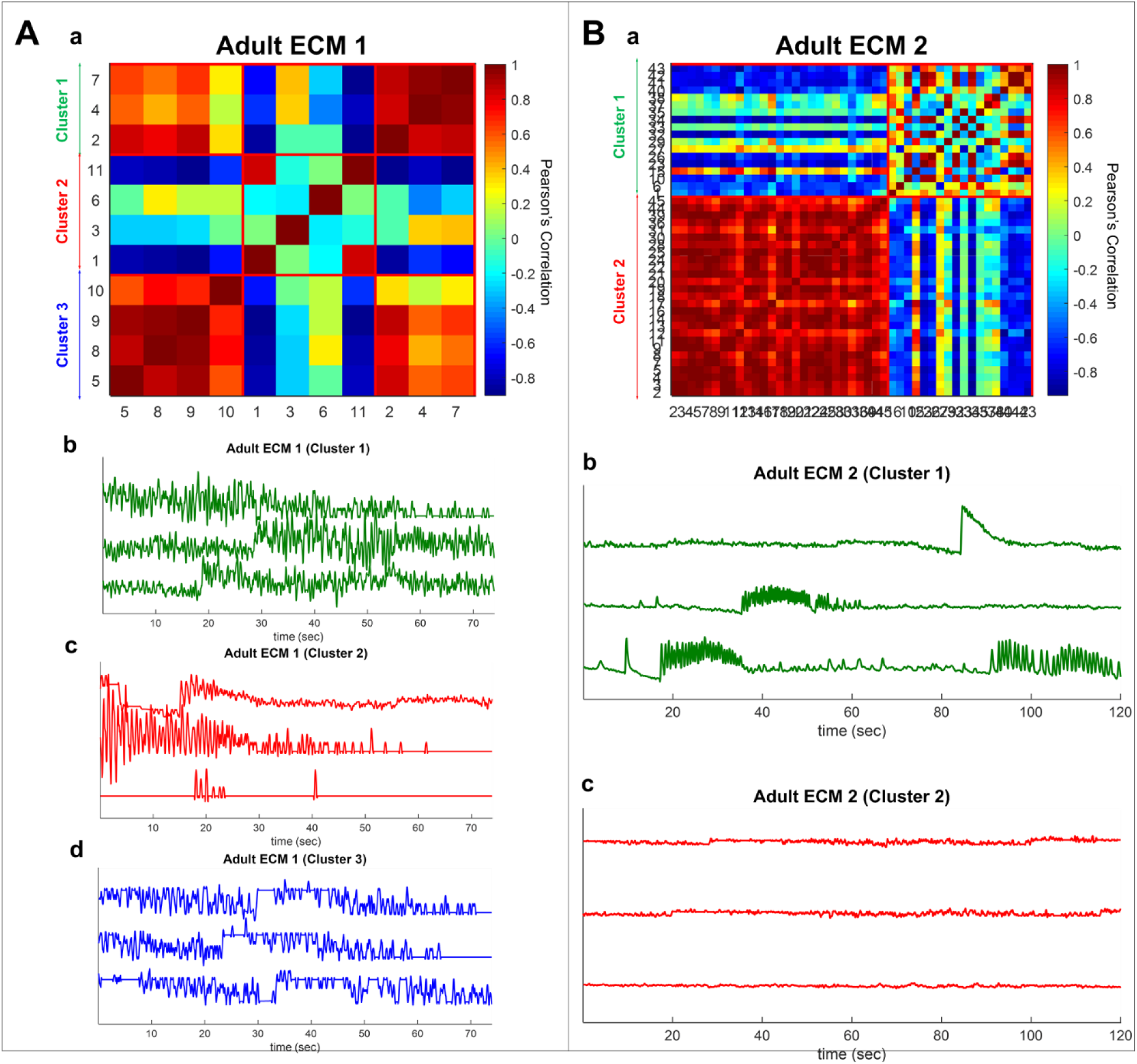
Spontaneous calcium activity in 3D cultures of differentiating human neural stem cells at 7 months; cells from adult ECM containing constructs. Tonic oscillatory activity is captured in Cluster 1 (panel A only); sporadic spiking activity is captured in Cluster 2 (panel A) and Cluster 1 (panel B); quiescent state is captured in Cluster 3 (panel A) and Cluster 2 (panel B).

**Supplementary Figure 8:**
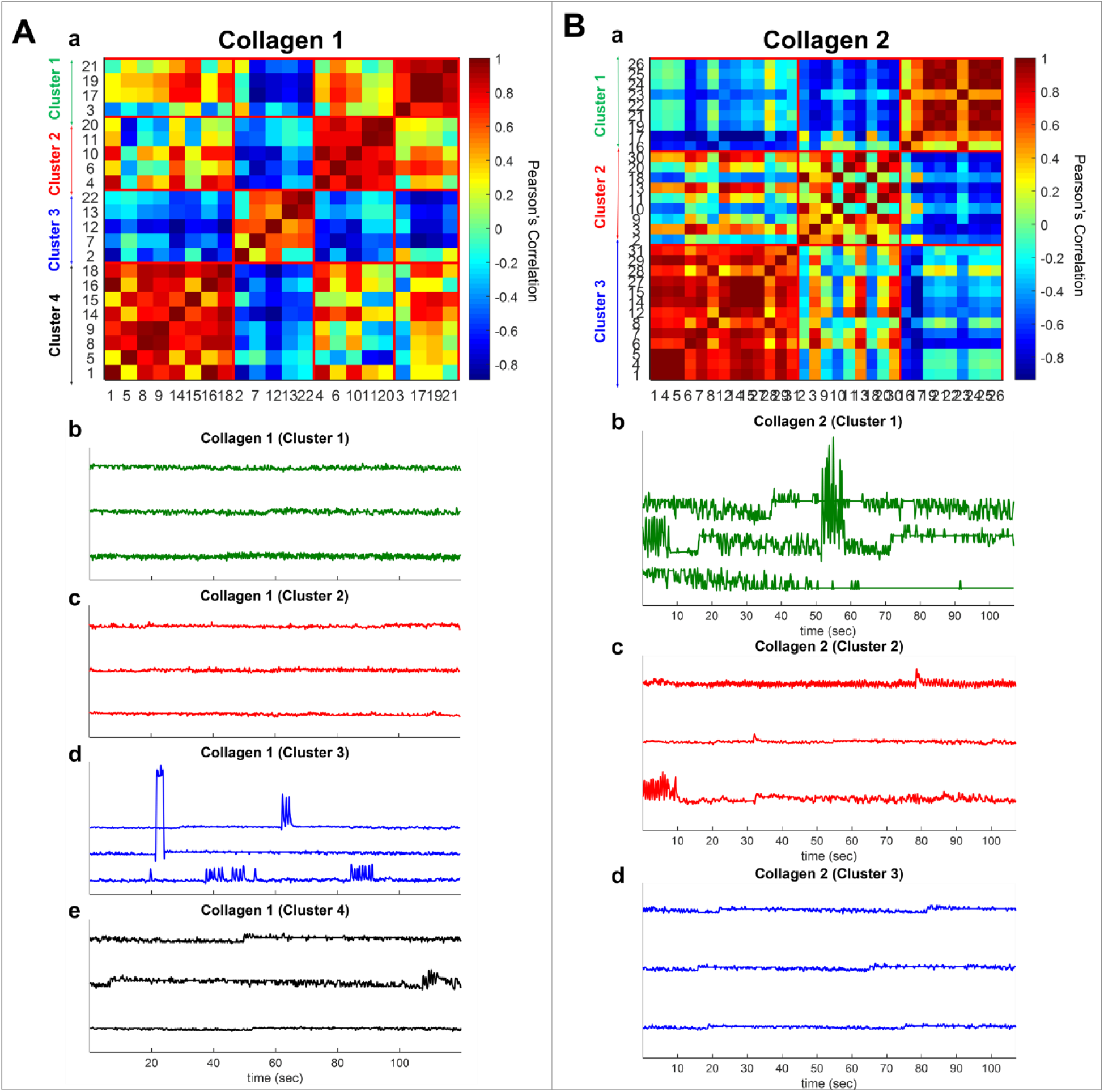
Spontaneous calcium activity in 3D cultures of differentiating human neural stem cells at 7 months; cells from unsupplemented constructs. Tonic oscillatory activity is captured in Cluster 1 (both panel A & B); sporadic spiking activity is captured in Cluster 3 (panel A) and Cluster 2 (panel B); quiescent state is captured in Cluster 2 & 4 (panel A) and Cluster 3 (panel B).

**Supplementary Figure 9:**
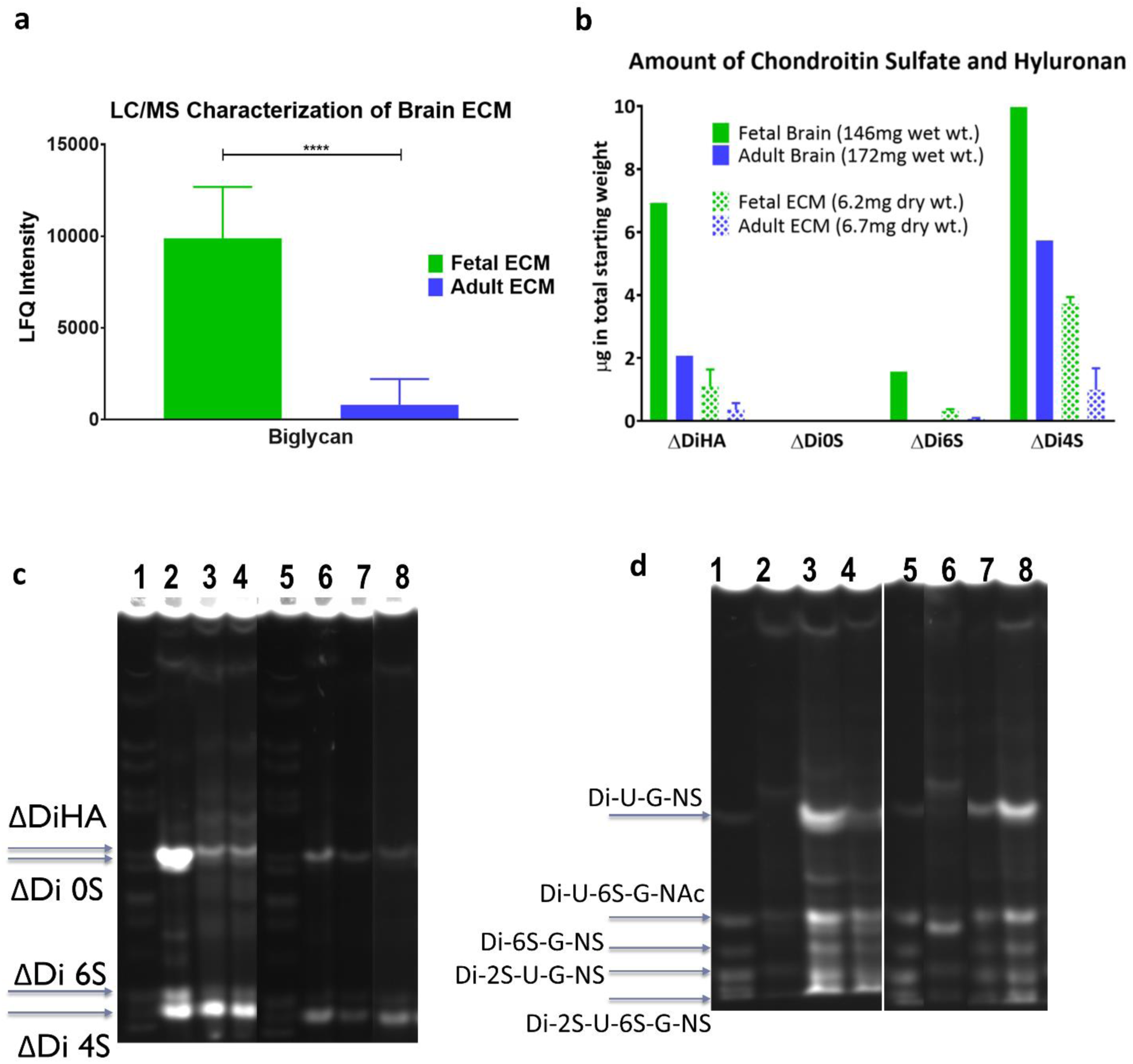
Characterization of decellularized porcine brain extracellular matrix. (a) LFQ intensities corresponding to biglycan fragments observed in fetal versus adult porcine brain decellularized matrix, following LC/MS analysis. Unpaired two-tailed t-test, n=3, p=0.0073. (b) Overall higher quantity of GAGs present in fetal brain, with corresponding higher retention in the extracted fetal brain ECM based on FACE analysis. (c) Characterization of decellularized fetal & adult brain ECM in comparison to fetal and adult whole brains by FACE. Chondroitin sulfate (CS) and hyaluronan (HA) bands in the fetal & adult porcine brain ECM. Lanes 1/5,2,3/4,6,7/8 correspond to standard, fetal brain, fetal ECM, adult brain and adult ECM, respectively. (d) Heparin sulfate (HS) bands in the fetal & adult porcine brain ECM over multiple extractions.

## Supplementary Videos

**Videos 1-1:** Spontaneous calcium activity in 3D cultures of differentiating human neural stem cells at 7 months; cells from a fetal ECM containing construct. Recorded at 20 frames per sec (fps), video shown at 50fps for 500 frames.

**Videos 1-2:** Spontaneous calcium activity in 3D cultures of differentiating human neural stem cells at 7 months; cells from an adult ECM containing construct. Recorded at 20 frames per sec (fps), video shown at 50fps for 500 frames.

**Videos 1-3:** Spontaneous calcium activity in 3D cultures of differentiating human neural stem cells at 7 months; cells from an unsupplemented construct. Recorded at 20 frames per sec (fps), video shown at 50fps for 500 frames.

**Videos 1-4:** Spontaneous calcium activity in 3D cultures of differentiating human neural stem cells at 3 months; cells from a fetal ECM containing construct. Recorded at 20 frames per sec (fps), video shown at 50fps for 500 frames.

**Videos 1-5:** Spontaneous calcium activity in 3D cultures of differentiating human neural stem cells at 3 months; cells from an adult ECM containing construct. Recorded at 20 frames per sec (fps), video shown at 50fps for 500 frames.

**Videos 1-6:** Spontaneous calcium activity in 3D cultures of differentiating human neural stem cells at 3 months; cells from an unsupplemented construct. Recorded at 20 frames per sec (fps), video shown at 50fps for 500 frames.

